# Niche Availability and Competitive Facilitation Control Proliferation of Bacterial Strains Intended for Soil Microbiome Interventions

**DOI:** 10.1101/2023.10.17.562719

**Authors:** Senka Čaušević, Manupriyam Dubey, Marian Morales, Guillem Salazar, Vladimir Sentchilo, Nicolas Carraro, Hans-Joachim Ruscheweyh, Shinichi Sunagawa, Jan Roelof van der Meer

## Abstract

Microbiome engineering, the rational manipulation of microbial communities and their habitats, is considered a crucial strategy to revert dysbiosis. However, the concept is in its infancy and lacks experimental support. Here we study the ecological factors controlling the proliferation of focal bacterial inoculants into taxa-complex soil communities and their impact on resident microbiota. We demonstrate using standardized soil microbiomes with different growth phases that the proliferation of typical soil inoculants depends on niche competition. By adding an artificial, inoculant selective niche to soil we improve inoculant proliferation and show by metatranscriptomics to give rise to a conjoint metabolic network in the soil microbiome. Furthermore, using random paired growth assays we demonstrate that, in addition to direct competition, inoculants lose competitiveness with soil bacteria because of metabolite sharing. Thus, the fate of inoculants in soil is controlled by niche availability and competitive facilitation, which may be manipulated by selective niche generation.

**Teaser:** Typical bacterial inoculants for soil microbiome engineering suffer from facilitating growth of native resident microorganisms

## INTRODUCTION

Microbiomes, the collective composite of microbial taxa and their habitats, play crucial roles in the functioning and health of their hosts or environments. Imbalanced or dysfunctional microbiomes pose a great challenge as they may present unstable developmental trajectories with a greater tendency for outgrowth of pathogens, reduced diversity, and/or diminished key ecological processes ^1−4^. Consequently, there is an important need to understand whether and how interventions can be directed to equilibrate a microbiome’s compositional or functional trajectory ^5,6^.

A classical intervention to alter microbiome composition is the inoculation of one or more microbial strains with specific functionalities ^7^, for example, to provide pollutant-degrading capacity for a contaminated site ^8,9^ or to enhance secondary metabolite production to deter potential plant pathogens ^10−12^. However, despite the conceptual simplicity, such inoculations mostly fail to produce the intended effect ^7,12,13^. The reasons for failure can be manifold but are reflected in the poor proliferation of the inoculated strain(s) within the targeted microbiome. Typically, probiotic therapies compensate for this effect by frequent (e.g., daily) reapplication of the strain mixture to temporally manifest and maintain the required function (14, 15). Nonetheless, the fundamental questions of why newly inoculated strains often struggle to establish in target microbiomes and, accordingly, why taxa-complexity provides microbiomes with invasion ‘resistance’ remain unresolved.

Niche availability is thought to be an important factor determining successful inoculant proliferation (16). The growth and development of a species-diverse microbiota is likely to exploit all carbon, nutrient, and spatial niches in their habitat leaving few open niches for incoming species to proliferate ^14−16^. Consequently, as a result of emerging functionalities by the microbiome the habitat conditions may change further ^17^ to disfavor easy access and opportunities for new strains to grow. Furthermore, cells of freshly inoculated strains may not find appropriate spatial niches to be protected against predation, e.g., by protists ^18^, resulting in their general decline ^19^, or fail to establish profitable interactions with resident microbiota species ^7^. Many of these arguments have not been subjected to systematic experimental testing of both the receiving microbiome and the introduced inoculant strain, and mechanistic concepts have been developed based on a small number of, frequently pathogenic, strains. We ourselves and others have recently argued how selective inoculation studies into defined microbiota, a concept we named N+1/N–1 engineering ^20^, can be used to uncover underlying mechanisms of community assembly and development to guide future intervention practices.

The major objectives here were to study the importance of potential niche availability and interspecific interactions on the proliferation of soil inoculants intended for use either to reinforce xenobiotic compound metabolism or to provide plant-growth beneficial functions. Four inoculants were selected: two are capable of degrading monoaromatic compounds such as toluene (*Pseudomonas veronii* 1YdBTEX2 and *Pseudomonas putida* F1) ^21,22^, one is a plant-beneficial bacterium (*Pseudomonas protegens* CHA0) ^23^, and one was selected as a non-soil strain control (*Escherichia coli*). To test the effects of niche availability we cultured standardized, taxonomically diverse, naturally-derived soil communities (NatComs) inside sterile soil microcosms according to a previously developed protocol (*27*). This system allowed us to test three conditions of niche availability for introduced inoculants. First, where all potential niches were assumed to be available and the inoculant would be in direct competition with NatCom to colonize the microcosm. Second, where the majority of niches were assumed to be occupied following precolonization by NatCom, after which the inoculant was introduced. Finally, we tested the effect of generating an inoculant-selective niche in the soil microcosms in the form of bioavailable toluene. Inoculant and NatCom populations were followed over time in their soil habitats, to estimate the realized niche from the extent of inoculant proliferation, and quantify any resulting changes in community diversity. To better understand the potential impact of biotic interactions on inoculant proliferation, we studied randomized paired-growth interactions between inoculant and soil bacteria in micro-agarose beads ^24^. Finally, by metatranscriptomic analysis of enriched expressed gene functions, we evaluated *in situ* metabolic interactions by resident bacteria in a broader variety of soils as a consequence of *P. veronii* growth and its metabolism of toluene. Our results clearly indicate the generally poor proliferation of soil inoculants is a result of limited niche availability and their tendency to lose productivity as a result of metabolite sharing with resident soil bacterial taxa. However, the provision of an inoculant specific niche improves inoculant survival and allows its functional integration into the resident microbiome network.

## RESULTS

### Producing taxa-diverse soil-cultured microbial communities in growing or stable states

To investigate the potential for inoculants to find colonizable ecological niches within a complex resident soil microbiota, we produced two standardized types of community physiological ‘states’: (i) a growing resident community (*GROWING*) and (ii) a steady-state resident community (*STABLE*). Our hypothesis was that the introduction of an inoculant simultaneously with a resident community into a niche replete soil habitat would give all strains equal opportunity to colonize available niches and proliferation would be increased, whereas in the case of a STABLE niche depleted resident community inoculants would find fewer available niches and proliferation would suffer.

Resident soil microbiota was grown from existing soil communities (NatComs) ^25^, which had been maintained for 1.5 years in soil microcosms. The soil microbiota were revived by mixing the colonized soil 1:10 (*v*/*v*) into freshly prepared soil microcosms. In total 50 microcosms were produced, of which 5 were used for community analysis in Phase 1 and the rest for Phase 2, see below. Microcosms all consisted of a sterilized silt matrix supplemented with soil organic carbon solution (Fig. 1A). Diluted NatComs were incubated for one month during which rapid growth was observed in the first days post dilution followed by a stabilization of the community size at 8×10^8^ cells g^−1^ soil (Fig. 1B). This density is comparable to the previously observed NatCom community size ^25^ and is similar to typical microbial cell densities in top soils ^26,27^, suggesting a maximum carrying capacity of the matrix and, thus, utilization of the available nutrient niches. Community succession was characterized by an initial increase in the most abundant phyla Firmicutes and Proteobacteria followed by slower-growing taxa, such as those from the Planctomycetes phyla (Fig. 1C). Lesser abundant members of Verrucomicrobia, Chloroflexi, Bacteroidetes, and Actinobacteria were also detected in the revived NatComs after one month (Fig. 1C). NatCom succession led to a temporary decrease in detectable richness and community evenness, which slowly increased and stabilized (Fig. 1B, *P*= 0.0556, Wilcoxon test comparison of different replicates from T_0_ to Day 23 and Fig. S1). After one month, the revived NatComs again resembled their starting material (Fig. 1D, PcoA ordination based on Unifrac distances at species level). *Inter alia*, this experiment also showed that cultured taxonomically-diverse soil communities can be maintained within the soil for extended time periods without extensive taxa loss.

**Figure 1.**
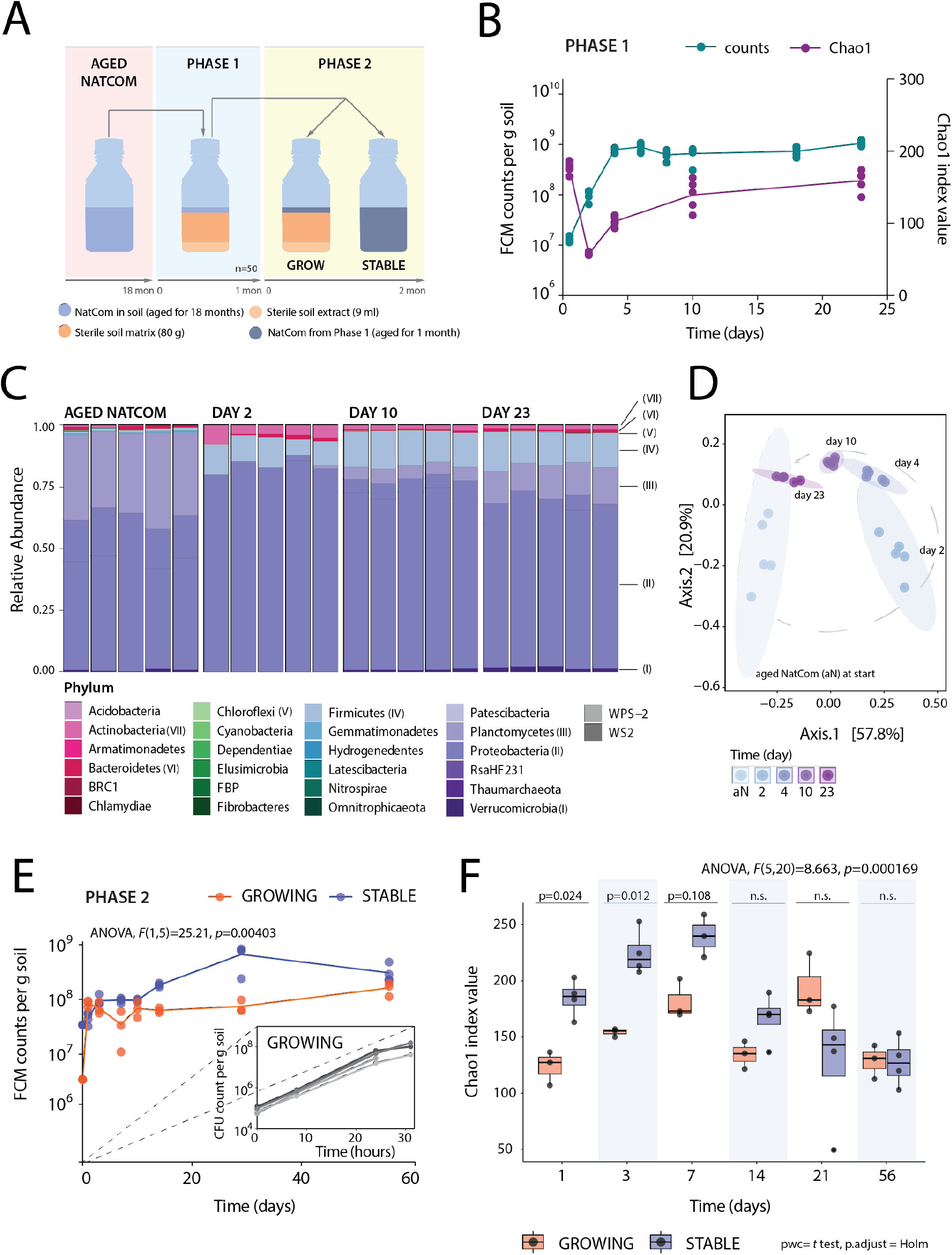
Producing standardized soil microbial communities for testing inoculant niche availability. **(A)** Experimental approach to obtain resident soil microbial communities (NatComs) in either a growing or stable state. In Phase 1, 1.5 year old, stored soil NatComs were revived by diluting material into fresh sterile soil microcosms and incubating for one month. After one month, pooled, revived NatComs were used for Phase 2, either directly (community condition ’STABLE‘) or diluted 1:10 (*v*/*v*) with fresh microcosm material (condition ’GROWING’). **(B)** Community growth flow cytometry (FCM) counts (left y-axis, cyan) and Chao1 index values for richness at amplicon sequence variant (ASV) level during Phase 1 (right y-axis, purple line). Dots show individual replicate measurements. Replicates are the same for all time points (repeated measures) except for T_0_, which is a sample of the total pool used for Phase 1 inoculation. **(C)** Phyla relative abundance changes during Phase 1. Individual stacked bars per time point show biological replicate compositions. The most abundant phyla are indicated with Roman numerals and described in the legend. **(D)** Trend in community development during Phase 1 (from light blue to dark magenta along the gray dashed line) represented on PCoA ordination of Unifrac distances (ASV level). Ellipses group replicates at each time point. **(E)** Mean Phase 2 community cell densities over time (FCM counts; dots indicating individual biological replicates). *P*- and *F*-values refer to cell density differences between GROWING and STABLE NatCom sizes (two-way repeated measures ranked ANOVA, S1 Dataset). Inset plot shows repeated experiment for community growth during the first 30 h upon dilution 1:100 (*v*/*v*) with fresh microcosm material quantified using colony forming units (CFU) per g of soil (shades of gray represent biological replicates). **(F)** Chao1 richness at ASV level in GROWING (orange) and STABLE (blue) NatComs during Phase 2. Boxplots show median and quartiles with individual values represented as black dots. Full *p*-values are indicated only if < 0.05 before or after *p*-value adjustment and otherwise considered non significant (n.s.). ANOVA refers to a repeated measures two-way test for the effect of community state and time interaction.

At the end of Phase 1 (Fig. 1A), all soil microcosms were pooled (the remaining 45, see above) and divided into two sets that served as resident background for testing inoculant proliferation. One part was diluted (1:10 *v*/*v*) in fresh soil matrix and nutrient to create a GROWING condition in Phase 2 (Fig. 1A), whereas the STABLE condition was produced purely from pooled and mixed material from the end of Phase 1 filled in new bottles (Fig. 1A). The GROWING NatCom showed rapid growth (19.8-fold average increase) in the first 10 days to a final average 47.2-fold size increase after 56 days (Fig. 1E, inset). Cell densities in the STABLE NatComs also increased, perhaps because of the pooling and mixing process at the end of Phase 1, but less than in the GROWING NatCom (2.8-fold after 10 days and 8.9-fold at day 56; Fig. 1E). During Phase 2, the GROWING NatCom cell densities remained lower than in the STABLE NatComs (Fig. 1E, *P*=0.00403, F(1,5)=25.21, two-way repeated measures ranked ANOVA) but eventually reached similar values at day 56 (*P*=0.1121, paired t-test). As expected for the faster proliferation of the GROWING NatComs during the first week, their taxa richness was initially lower than that of STABLE NatComs, but became similar from Day 14 onwards (Fig. 1F). This apparent lower diversity is due to the saturation of sample sequencing depth by faster-growing species and a subsequent reduction in the detectable species count. Over time slower-growing species become more abundant and are redetected. The Shannon diversity index of GROWING NatComs also remained slightly lower throughout the incubation than that of STABLE NatComs (Fig. S1, ASV-levels; *P* = 0.00014). We thus concluded that, while the dynamic succession of GROWING and STABLE NatComs varied, the equivalent richness and size of either community meant they were suitable for testing the effect on inoculant proliferation.

### Inoculant establishment is dependent on niche availability and inoculant characteristics

We tested the four selected bacterial inoculants (*P. putida*, *P. protegens*, *P. veronii*, and *E. coli*) for their potential to proliferate in microcosms under different conditions: axenically (ALONE), concomitantly with the freshly diluted NatCom (Phase 2 GROWING, starting at t=0 as in Fig. 1E), or after NatCom establishment (STABLE, also starting at t=0 in Fig. 1E). All inoculants constitutively expressed an introduced genetically encoded mCherry tag, which facilitated the quantification of their specific population size. Axenically, all three pseudomonads reached similar cell densities (1–3×10^7^ cells g^−1^ soil) in the microcosms after 3 days of incubation (Fig. 2A, ALONE), which corresponds to a 100-fold or more increase compared to their inoculated population sizes (1×10^5^ cells g^−1^ soil). This demonstrated that the strains could grow at the expense of available resources within the soil extract. As expected, *E. coli* proliferated poorly and only increased its cell density by 10–12-fold within the soil (3–4 generations; Fig. 2A, ALONE). Over time all axenic populations slowly decreased in size suggesting some cell death occurred.

**Figure 2.**
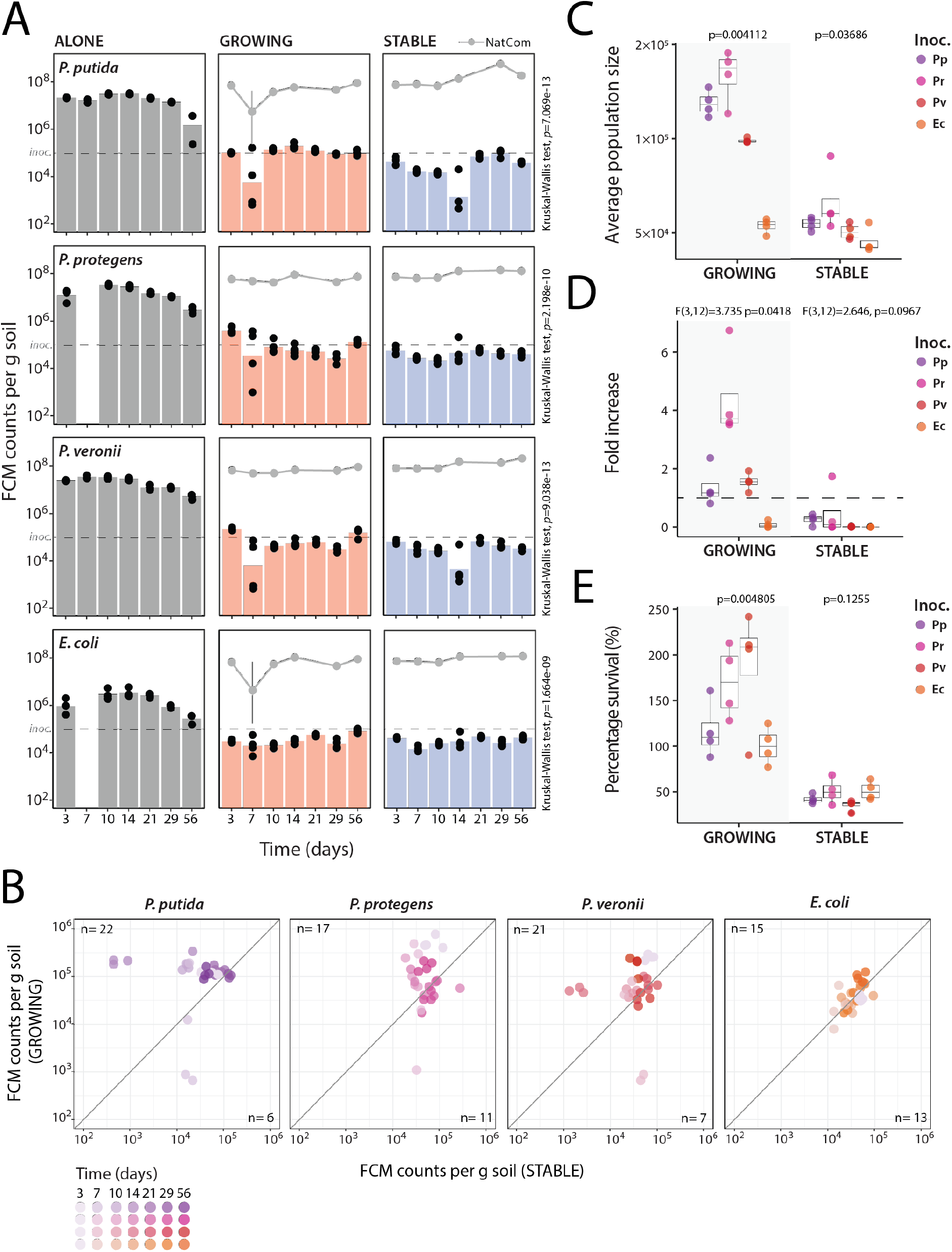
Dependence of inoculant establishment on the growing state of resident communities. **(A)** Bars show mean inoculant (*P. putida, P. veronii, P. protegens,* or *E. coli*) population sizes at each time point in sterile microcosms (ALONE, gray) and microcosms with GROWING (orange) or STABLE NatComs (blue). Black circles indicate individual biological replicate values (*n =* 4). Population sizes are expressed as FCM cell counts per g of soil. Gray lines in GROWING or STABLE subplots connect the mean total community size measurements in the same samples (vertical lines representing ± one standard deviation of the mean). *P*- value of Kruskal-Wallis test is indicated on the left of each group of subplots. Dashed lines indicate the calculated inoculum population size. **(B)** Differences of inoculant population sizes in GROWING versus STABLE resident communities. Dots represent individual replicate inoculant population size measurements from GROWING or STABLE NatComs paired by the same incubation time point, and colored by inoculant from GROWING or STABLE series (purple for *P. putida* (Pp), pink for *P. protegens* (Pr), red for *P. veronii* (Pv), and orange for *E.coli* (Ec), and color gradients follow time points, as in the scale. The diagonal indicates the *null* hypothesis of no difference between inoculant survival in growing or steady-state communities. The number of dots above and below the diagonal is signalled on the plot as value *n*. **(C)** Average inoculant population size in GROWING (grey background) or STABLE NatComs during the entire experiment (i.e., mean of all sampling time points, except T_0_). Boxplots show median, quartiles, and individual values; dot colors as in (B). *P*-values refer to differences among inoculants by separate Kruskal-Wallis testing for GROWING and STABLE NatComs. **(D)** As for (B), but for the maximum observed fold-difference of inoculant density compared to T_0_. Dashed lines indicate a fold-difference of 1. *P*- and *F*-values refer to one-way ANOVA test results for difference among inoculants. **(E)** Percent inoculant survival after two months (as the ratio of inoculant population size after two months divided by the initial inoculum size). *P*-values as in (C).

In contrast, when co-inoculated with GROWING NatComs or inoculated into STABLE NatComs all inoculant populations attained significantly lower population sizes than axenically in soil microcosms (Fig. 2A, Kruskal-Wallis test, *post hoc* Dunn pairwise test, S1 data). The average inoculant population sizes here reached between 5×10^4^–2×10^5^ cells g^−1^ soil, depending on the NatCom state and inoculant (Fig. 2B and C), but remained relatively stable until the end of the experiment (Fig. 2A, approx. two months). The growth and survival of inoculated pseudomonads was better in GROWING than in STABLE NatComs (Fig. 2B and C), whereas the population sizes of *E. coli* were the lowest among all inoculants and no different in GROWING or STABLE NatComs (Fig. 2B; 2C, Wilcox signed-rank test, *P*=0.68). Among the pseudomonads, *P. protegens* showed the highest net population expansion (in comparison to the inoculated level; Fig. 2D), whereas both *P. protegens* and *P. putida* showed the highest average relative abundances (Fig. 2C; Fig. S2). The population densities of all pseudomonads after two months demonstrated they had survived in GROWING NatComs and maintained a size higher than their initial inoculum (Fig. 2E). These results support our hypothesis that the soil inoculants (all pseudomonads but not *E. coli*) were able to find more available niches for their establishment within a diverse soil resident community under GROWING conditions than in the background of an established STABLE community. The difference in inoculant proliferation in axenic microcosms compared with community growth indicated that only around 1% of the potential nutrient niche for the (pseudomonad) inoculants is available within a taxonomically diverse resident soil community (Fig. 2A, ALONE *vs.* GROWING or STABLE), thereby indicating that niche competition is a major factor that limits their expansion. Furthermore, these results showed that pseudomonads have better colonization success in soil than a poorly soil-adapted strains such as *E. coli*. However, they did not attain cell densities higher than two times the inoculum size.

### Creation of a specific niche favors inoculant establishment in resident communities

To test whether inoculant outgrowth is limited by niche competition and not by *a priori* predation, we exploited the capacity of one of the inoculants (*P. veronii*) to metabolize toluene, which we could add as a unique additional carbon substrate (assuming that the ability of NatCom strains to metabolize toluene toluene would be limited). The GROWING and STABLE NatComs were thus exposed to toluene, which was provided in the gas phase of the microcosm from where it could reach the cells in the soil pore aqueous phase by diffusion.

Supplementation of toluene had no statistically significant effect on the sizes of the STABLE NatCom (Fig. 3A, *P*_STABLE_=0.583) but significantly increased their sizes by an average of 1.5-fold in GROWING NatComs (*P*_GROWING_= 0.00151). In contrast, *P. veronii* attained 100–200-fold larger population sizes in the presence of toluene compared to unamended microcosms, irrespective of being co-inoculated with GROWING or inoculated into STABLE NatComs (Fig. 3B, Kruskal-Wallis *P*=2.2×10^-16^, *post hoc* Dunn test; S1 data). Eventually, the *P. veronii* populations declined in the presence of toluene but still maintained significantly higher levels than in its absence (Fig. 3B). This experiment thus demonstrated that the proliferation of an inoculant is significantly improved when it finds a specific and selective niche. It also suggested that it is effectively the absence of a selective niche and competition for shared niches that limited its development in the unamended NatCom microcosms.

**Figure 3.**
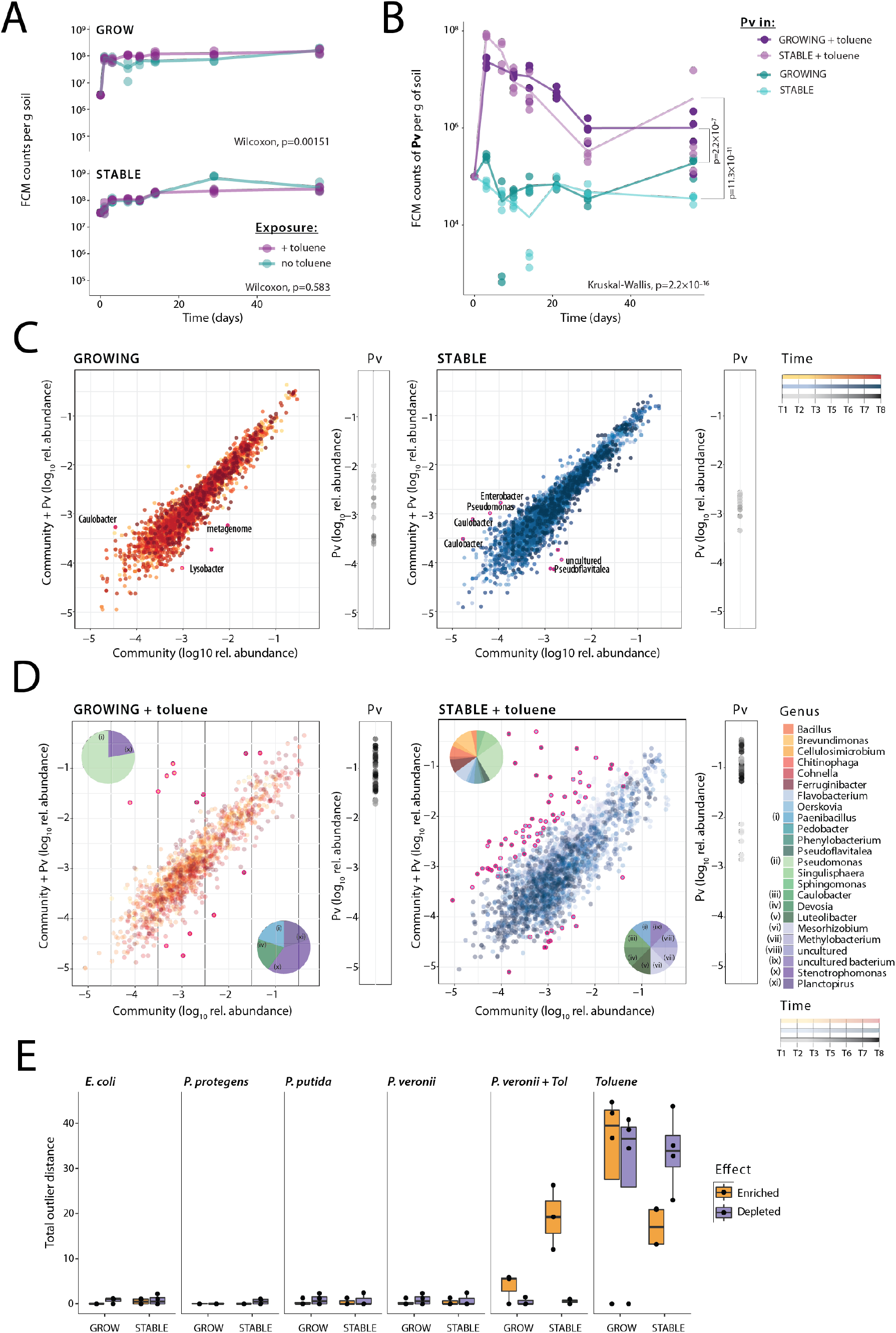
Effect of a toluene selective nutrient niche on growth and survival of *P. veronii* within resident soil communities. **(A)** Community sizes over time in presence or absence of toluene but without inoculant. Dots show replicate FCM cell counts per g soil (*n* = 4 separate microcosms) with lines connecting the mean values. *P*-values result from Wilcoxon signed rank testing, comparing the effect of toluene on the total sizes of either GROWING or STABLE communities. **(B)** *P. veronii* population development within GROWING or STABLE NatCom exposed to toluene (shades of pink) or not (shades of green). Population data from *P. veronii* without toluene are reproduced from Fig. 2A for clarity. *P*-value from Kruskal-Wallis test for the difference between toluene- or non-treated samples. *post hoc* Dunn test *p*-values (with Holm’s adjustment) refer to differences of *P. veronii* population sizes at day 56 as a function of toluene treatment between GROWING and STABLE NatComs. **(C)** and **(D)** Deviations of operationally-defined taxonomic unit (OTU level, genus name indicated) abundances in *P. veronii* (Pv) inoculated microcosms, without (D) or with toluene (E). Dots represent time-paired log_10_-transformed relative abundances of the same OTU across all biological replicates (arbitrarily paired among replicates) and treatments, as indicated. Upward deviations from the diagonal line indicate taxa enrichment in the inoculated microcosms, whereas downward deviations indicate taxa depletions. Magenta circles emphasize differences of more than one log to the expected value (i.e., the diagonal). Relative abundances of *P. veronii* (Pv) are presented within separate subplots on the side (color gradients follow time points as in the scale). Pie charts in (D) show relative abundances of all deviated taxa per condition (additionally labelled with roman numerals for clarity). **(E)** Total outlier distance, per treatment or inoculant, of all taxa more than ten-fold enriched (in yellow) or depleted (in purple) compared to non-inoculated or non-treated controls.

Considering that in toluene amended microcosms *P. veronii* composed 10%–20% of the total community size, we hypothesized that this may have caused secondary effects on resident populations. We thus compared paired taxa abundances in the absence or presence of *P. veronii*, per treatment, and over time (e.g., Fig. 3C and D). In the absence of toluene but with inoculated *P. veronii* there were only very few taxa outliers (defined as having 10-fold higher or lower abundances than expected for equal proportions), in either GROWING or STABLE NatComs (Fig. 3C). Outliers concerned a variety of low-abundance taxa, such as Caulobacter, Enterobacter, Lysobacter, Pseudomonas, and Pseudoflavitea (Fig. 3C), but appeared spurious as did not occur reproducibly across replicates and at more than one time point. The absence of clear effects was not surprising given the relatively low attained population size of *P. veronii* in these microcosms (<1%; Fig. 3C, Pv subplots).

In contrast, more dramatic shifts in taxa abundances were observed in presence of toluene and, perhaps counterintuitively, more in STABLE than GROWING NatCom (Fig. 3D). GROWING NatCom exposed to toluene and inoculated with *P. veronii* were notably depleted in Planctopirus, Devosia, and Paenibacillus taxa, enriched for Pseudomonas, and both depleted and enriched in various Stenotrophomonas strains (Fig. 3C). Inoculation of *P. veronii* into STABLE NatComs led primarily to the enrichment of a variety of taxa and, to a lesser extent, the depletion of others (Fig. 3D). Across all conditions in the absence of toluene, the difference in magnitude of secondary taxa changes (quantified as the total outlier distance) was almost undetectable for any of the inoculants (Fig. 3E). In contrast, taxa changes were more pronounced in the case of *P. veronii* inoculation in the presence of toluene, and largest for exposure of NatComs to toluene without *P. veronii* inoculation (Fig. 3E, Fig. S3). Interestingly, the *P. veronii* population size exposed to toluene declined less rapidly when resident NatCom was present (irrespective of GROWING or STABLE condition), compared to when it was growing axenically in microcosms (Fig. S3). This indicated, therefore, that *P. veronii* inoculation not only alleviates the negative effects of toluene exposure on the NatComs but that its longer-term survival benefited from the resident community.

### Inoculants lose productivity but favor growth of soil community members in random paired assays

To gain a better understanding of the interactions between introduced inoculants and existing resident communities, we transitioned from system-level experiments to exploring a multitude of potential pairwise interactions between our chosen inoculants and resident soil community members. For this purpose, we employed a method where single inoculant cells are randomly encapsulated and incubated with isolated soil cells within 40–70 µm agarose beads ^24^. In contrast to the work above with standardized NatComs, we here used bacterial cells freshly washed from their natural soil matrix, thereby expanding the range of taxa- inoculant combinations being explored (Fig. S4). We hypothesized that, because of the proximity of founder cell pairs, growth interactions would lead to deviations in the average microcolony size distribution of inoculant or soil resident compared with either member growing individually.

Paired growth was quantified by estimating the size of fluorescent microcolonies inside beads at different incubation times. Inoculant colonies were distinguished from the fluorescently stained soil taxa courtesy of their mCherry-fluorescence labels. The average size of encapsulated *P. veronii* microcolonies, incubated with soil extract as the sole carbon and nutrient source, increased more over time if *P. veronii* was incubated alone (Fig. 4A, PV ALONE) compared to beads where *P. veronii* was paired together with soil community (Fig. 4A, PV WITH SC, p=0.0005, Fisher’s two-tailed distribution test). Inversely, soil cells appeared to benefit from incubation with *P. veronii*, as the average microcolony size of soil cells increased when *P. veronii* was present as a partner (Fig. 4A, SC ALONE vs. SC WITH PV). This was not a result of differences in medium conditions because incidental beads in the inoculant-partner incubations with either only *P. veronii* (Fig. 4A, PV ALONE IN MIX) or only soil cells (Fig. 4A, SC ALONE IN MIX) showed similar average growth as the separate control incubations (incidental beads with individual taxa occupancy occur due to the random Poisson distribution of the encapsulation process).

**Figure 4.**
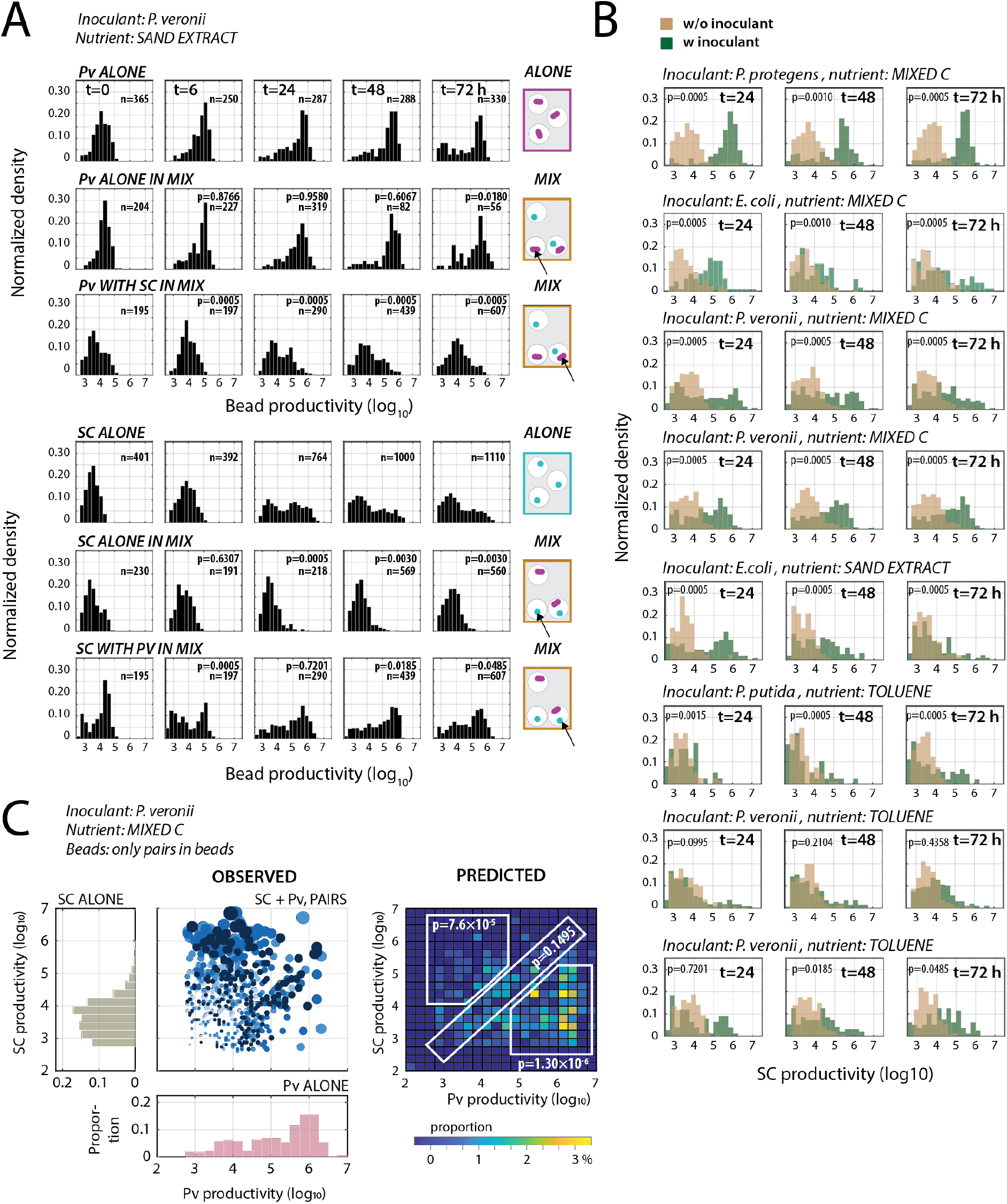
Paired productivities of inoculants with random resident soil cells. **(A)** Growth of either bead-encapsulated *P. veronii* (Pv) or soil cells (SC) alone, or of paired mixtures (e.g., Pv with SC, 1–2 random founder cells at start) in soil extract. Note the illustrations on the right explaining how mixed beads can by chance have true pairs of inoculant and soil cells or contain single cells of either Pv or SC. Plots show normalized histograms of microcolony sizes imaged from epifluorescence microscopy, expressed as the log_10_-value of the Syto9-observed pixel area × its mean fluorescence intensity. n, number of analyzed beads. *P*-values from Fisher’s exact test on distribution differences. **(B)** As for (A) but for different inoculants and media conditions and only showing the comparison of soil cells alone (in beads in the mixture, light brown) and in beads with soil cells and inoculants (dark green). P-values from Fisher’s exact test. Similar labels correspond to independent experiments. **(C)** Paired productivity plot of beads with only a single *P. veronii* and a single soil cell microcolony (OBSERVED), versus the distribution of bead-growth of *P. veronii* (Pv, salmon) or soil cells alone (SC). Pv and SC alone summed from time points 24, 48, and 72 h. Circles are proportional to the sum of the measured microcolony sizes (from light to dark blue correspond to time points 0, 6, 24, 48 and 72 h). The heatmap (PREDICTED) shows the expected paired bead summed productivities (in percentage, as per the color scale) from the individual measured microcolony sizes (i.e., Pv and SC ALONE) for the same number of beads as analyzed by microscopy. *P*-values correspond to the two-tailed t-test comparison of the variation of the total measured paired productivities inside the three regions (*n* = 3; 24, 48, and 72 h) with that in the simulations (*n* = 5). The upper left region shows higher SC productivity than expected, the lower right region shows lower inoculant productivity than expected, and the diagonal shows the same productivity for both microcolonies in a pair.

The same pattern was observed under most tested nutrient conditions (i.e., soil extract, a defined solution of 16 carbon substrates, and toluene) and with each of the four inoculants (Fig. 4B). This suggests that all the inoculants transform primary substrates into metabolites or otherwise increase local nutrient availability, leading to growth benefit of other soil cells in proximity (i.e., within the same bead). The process is disadvantageous for the inoculant itself as it reduces its own productivity. To show this more clearly, we selected beads from different time points containing exactly one inoculant and one soil cell taxon microcolony (Fig. 4C). This enabled us to detect shifts in paired productivities compared to the productivities expected from growth distributions of each member alone if they were indifferent to each other (Fig. 4C, *PREDICTED*). The experimental results clearly show a stronger than expected growth of soil taxa and consequently reduced growth of *P. veronii* in paired growth tests (p=7.6×10^−5^, two-tailed t-test, *n*= 3 experimental and 5 simulation replicates). We consistently observed similar outcomes across all four inoculant strains and in each growth condition, indicating the higher-than-expected growth of soil cells in paired beads with inoculants (Fig. S5). Thus, these findings demonstrated that, on average, all inoculants lose in substrate competition when in proximity of a soil cell, from which the latter can not only profit but more so than if growing alone. This also indicates that it is not only the direct loss of available niches, e.g. by faster-growing taxa, that limits inoculant growth but also the loss of competitiveness during niche transformation (i.e., *competitive facilitation*). Both effects help to explain why the inoculants established much more poorly in soil microcosms co-inoculated or precolonized with NatComs compared with sterile microcosms (Fig. 2A, GROWING and STABLE vs. ALONE).

To better understand whether competitive facilitation is soil taxa specific or general, we analyzed changes in OTU relative abundances as a function of growth conditions and inoculant using 16S rRNA gene amplicon sequencing of bead mixtures with or without inoculants (focusing only on *P. veronii* and *E. coli* as a negative control; Fig. 5). DNA isolated from beads after 48 h was dominated by 15–50 families (*n*=3, threshold > 10 reads). Notably, soil extract as a sole nutrient source enabled the highest taxa diversity and growth (Fig. 5A). As seen in the NatCom microcosms toluene caused a very selective effect and repressed the growth of most taxa. The number of families with significantly increased relative abundances in the presence of an inoculant was highest for *P. veronii* and toluene (eight enriched families and one depleted; adjusted P-value < 0.05) and lowest for *E. coli* and mixed-carbon substrates (Fig. 5A). The family of Micrococcaceae was enriched in all conditions when inoculated with *P. veronii* but not with *E. coli* (Fig. 5A, **M**). In comparison, when grown on the same carbon substrate mixtures, Micrococcaceae and Rhizobiaceae were more abundant in inoculations with *E. coli* and Burkholderiaceae and Enterobacteriaceae were more abundant with *P. veronii*, suggesting potential favorable metabolic interactions from inoculant to taxa from those families (Fig. 5B). Closer examination of Pseudomonadaceae (due to their generally high relative abundances) showed very little taxa sensitivity to the presence of the inoculant (except sequence variants ASVs I and II in Fig. 5C, which were more abundant with soil extract and absent on toluene) but did demonstrate sensitivity to the abiotic condition. For example, ASVs III and VI were relatively abundant in all incubations, whereas ASV-V was abundant in the presence of toluene but depleted in other substrate conditions (Fig. 5C). In contrast, ASV-IV was abundant in all conditions except for with toluene. Notably, these observations are based on relative normalized sequence abundances from incubations that contain mixtures of beads with pairs of inoculants and soil cells, as well as soil cells or inoculant cells alone (as illustrated in Fig. 4A). Thus, these results indicated that both inoculants stimulated soil taxa when growing in proximity (Fig. 4B), without being particularly selective at family (Fig. 5A) or even strain level (ASVs, Fig. 5C). Such evidence underscores the notion that successful inoculant proliferation in soil is challenging when only general substrates are provided due to competitive loss or, depending on the perspective, carbon facilitation to others.

**Figure 5.**
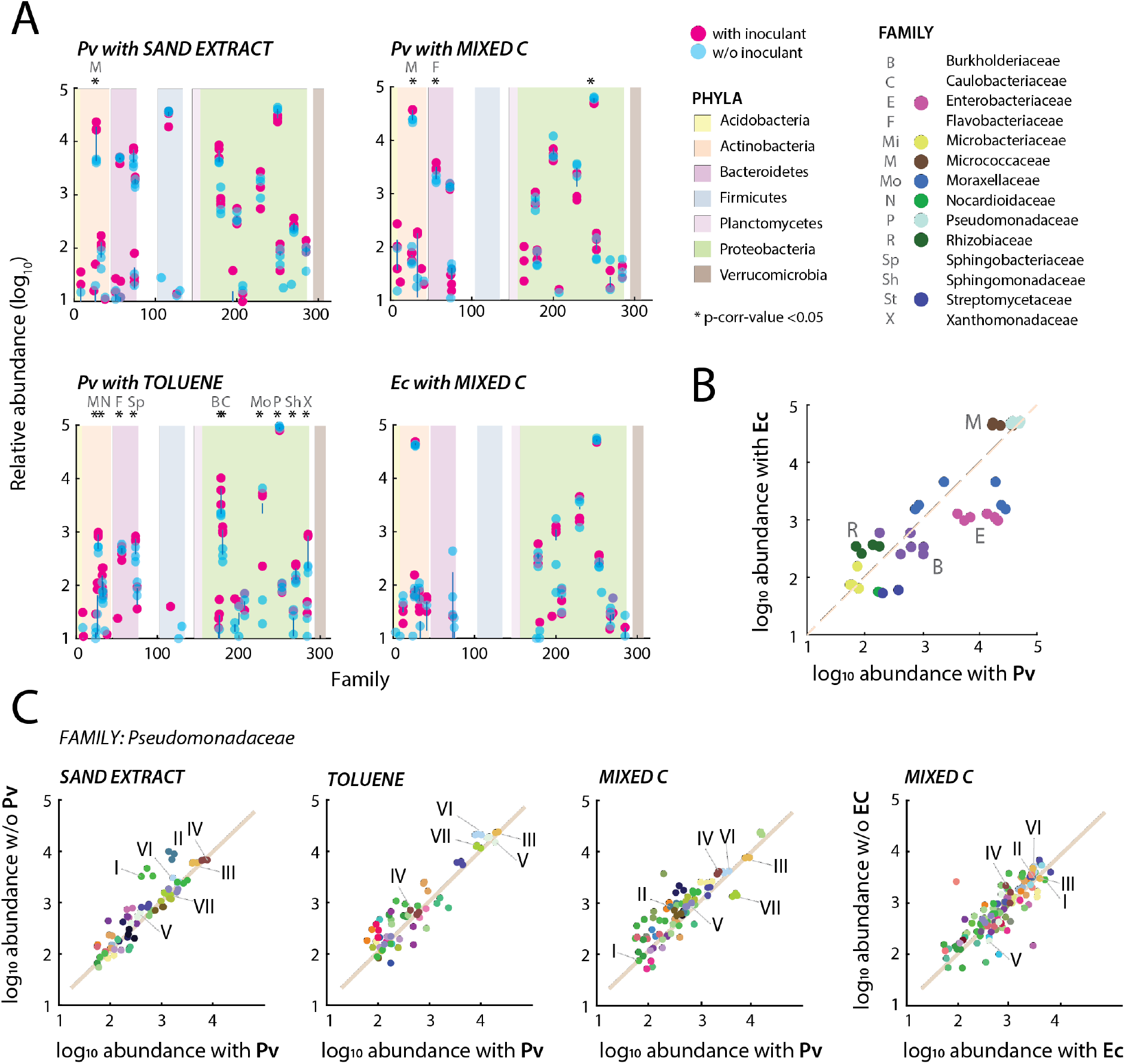
Diversity changes in inoculant-soil cell bead-encapsulated communities as a function of growth condition. **(A)** Log_10_ read abundances of family level summed taxa in bead-encapsulated communities after 48 h, either in the presence (magenta) or absence (blue) of an inoculant (resampled normalized abundances after removal of the inoculant reads). Dots indicate individual biological replicate values with a line connecting the means for the same taxa in absence or presence of the inoculant. Growth conditions as indicated. Background colors show phyla attribution. Asterisks denote significant differences (adjusted P-value < 0.05) with letters explaining the corresponding family. **(B)** Comparisons of soil-paired encapsulations with *P. veronii* versus *E. coli* incubated with Mixed-C (16 substrates), highlighting the names of consistently selectively responding families (legend in A). **(C)** Relative abundance changes of ASVs within the family of Pseudomonadaceae (after removal of the inoculant reads), as a function of incubation condition and inoculant. Dots show individual replicate values (paired arbitrarily between conditions with or without inoculant) with colors matching ASVs across conditions (specific examples emphasized with Roman numerals).

### Inoculant toluene metabolism triggers a variety of cross-feeding pathways in the resident community

Given the evident interactions taking place in the beads, we tried to delineate the potential cross-feeding network among resident soil bacteria arising from inoculation. We focused here specifically on inoculation of *P. veronii* and exposure to toluene, as under these conditions the inoculant population could establish sufficiently such that potential effects on resident bacteria might be detected. We hypothesized that while toluene provided a specific growth advantage to *P. veronii* within the NatComs, its metabolism could indirectly facilitate the growth of other taxa, as suggested by both the encapsulation experiments (Fig. 4C and Fig. 5A) and microcosm studies (Fig. 3D). Here, we took advantage of a previously conducted study where *P. veronii* was inoculated into two types of natural soils, Silt and Clay, and into a contaminated control soil from a former gasification site (Jonction) ^28^. The use of varied, non-sterile soils and materials beyond standardized microcosms is important here to demonstrate the more general nature of *P. veronii* inoculation successes and its impacts.

Total RNA isolated from microcosms containing each soil type at early and late time points (Fig. 6A) was subjected to metatranscriptomic sequencing, assembly, and annotation to quantify gene expression levels of the native soil taxa. *P. veronii* established in Silt and Jonction soils exposed to toluene, while Clay soils demonstrated higher resistance to inoculant establishment despite toluene addition (Fig. 6A). Soils where *P. veronii* was actively growing (Fig. 6A, Silt and Jonction) also showed higher abundances of transcripts for ribosomal proteins, which indicates increased activity and growth of the resident community (Fig. 6B). Notably, uninoculated but toluene-exposed Silt resident communities had higher abundances of transcripts for ribosomal proteins but only later in the incubation. Resident microbiota transcripts associated with aromatic compound metabolism were enriched under toluene exposure in presence of inoculated *P. veronii*, particularly for pathways linked to known metabolites of the *P. veronii* toluene degradation pathway (Fig. 6C and D; I, II and III; Fig. S6). Such enrichments were particularly prevalent in Silt microcosms compared to their corresponding inoculated toluene-free or inoculated and toluene- exposed controls (Fig. 6D). Toluene exposure in the absence of *P. veronii* also provoked an increase in transcript abundance of aromatic compound degradation pathways but generally at the later sampling point, suggesting some growth of native toluene degrading strains. Jonction, as expected for an already contaminated soil, carried high transcript levels of a higher functional diversity of genes for aromatic compound metabolism (Fig. S6 and S7). These transcripts could be assigned to close relatives of *Immundisolibacter cernigliae* and *Rugosibacter aromaticivorans* (Fig. S8), two known degraders of aromatic compounds ^29,30^. Transcripts related to aromatic compound metabolism were generally low in the Clay microcosms, probably because *P. veronii* did not proliferate well and no secondary effects had taken place (Fig. 6A). Interestingly, the sum of transcripts in the resident soil microbiota for the exploitable toluene degradation products followed a log-linear correlation with their overall growth state, as estimated from transcripts for ribosomal proteins (Fig. 6B). For a small number of increased aromatic compound metabolism transcripts we traced the potential source organism (Fig. S7). As expected, identified taxa were already enriched in Jonction but other stimulated taxa in Silt and Clay became enriched following toluene and *P. veronii* exposure in agreement with the observed NatCom stimulation (Fig. 3D). In summary, these results show that the metabolism of toluene by *P. veronii* can elicit a cascade of cross-feeding pathways among resident soil bacteria.

**Figure 6.**
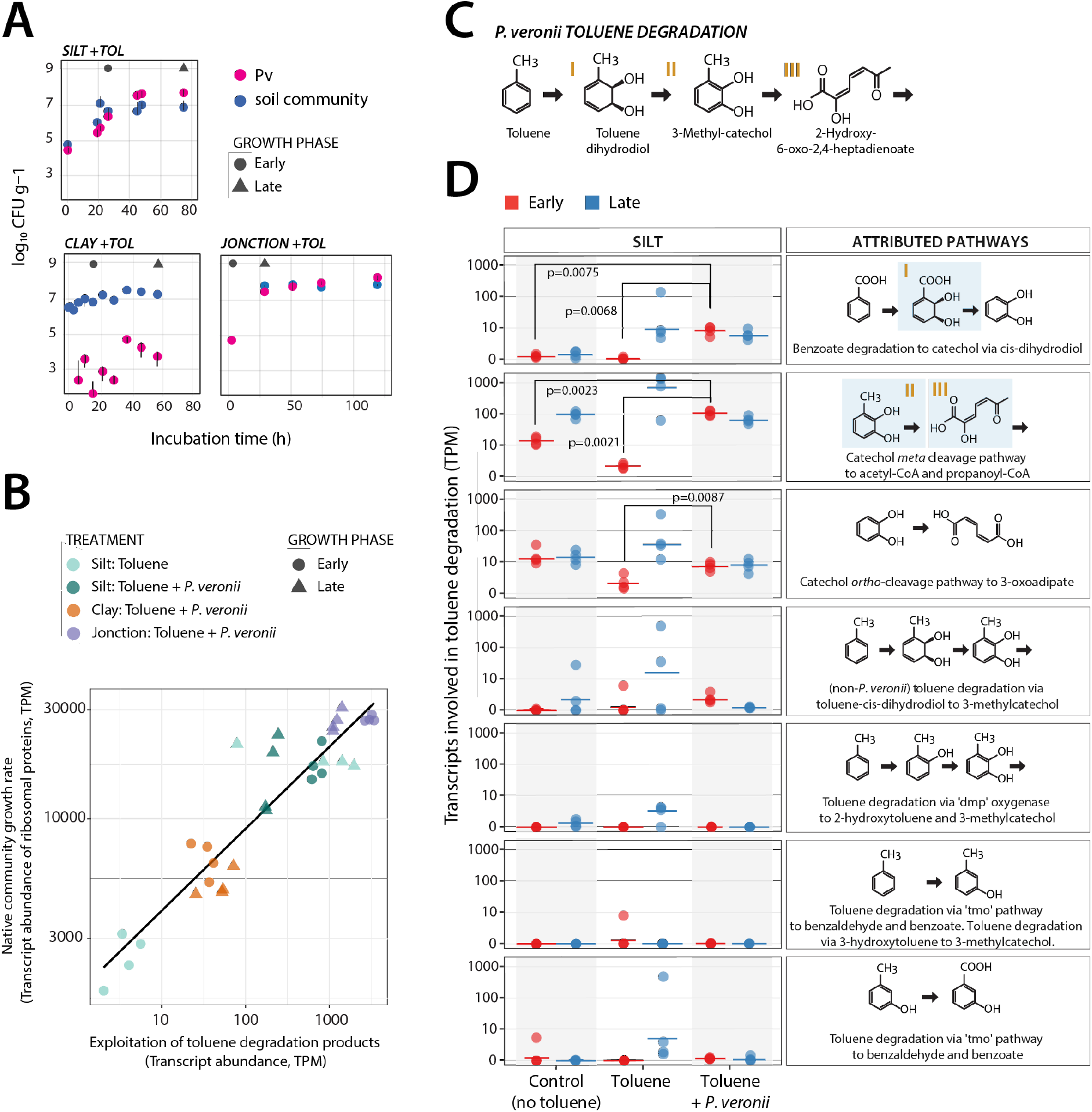
Exploitation of toluene degradation products from *P. veronii* by soil microbiota. (**A**) Survival or proliferation of inoculated *P. veronii* in Silt, Clay, or *Jonction* soils exposed to gaseous toluene (data replotted from Ref. ^28^). Data points are the mean from triplicate measurements of colony forming units of *P. veronii* (magenta) or resident soil microbiota (blue). Gray circle and triangle indicate time points for sampling of total community RNA (early and late time point, respectively). (**B**) Transcript abundances (transcripts per kilobase million (TPM) without *P. veronii* transcripts) of ribosomal proteins in the communities versus those of functions attributed to utilization of toluene or its metabolites (see panels C and D). (**C**) *P. veronii* toluene degradation pathway and major metabolic intermediates. Roman numerals correspond to highlighted intermediates in panel D. (**D**) TPM-values of transcripts annotated to the aromatic compound metabolic steps on the right for Silt in three conditions tested at two stages (*early* or *late*, as in panel A). Data points show individual values from quadruplicate experiments and a line indicating the median. *P*-values correspond to t-test statistics comparison of indicated sample replicate measurements (only shown if <0.05). For details of KEGG pathway attributions, see Fig. S6.

## DISCUSSION

Microbiome interventions based on strain inoculations are frequently frustrated by the poor proliferation of the inoculant and, thus, an insufficient display of their intended function ^7,12,13^. To better understand the underlying ecological conditions and mechanisms leading to poor inoculant proliferation, we systematically studied the potential of available niches on the establishment of a variety of inoculant strains in soil microcosms containing taxa-diverse resident soil microbiota. Our studies benefitted from the reproducible culturing of taxa-complex soil microbiomes, which enabled us to contrast the proliferation of inoculants concomitant to growing soil microbiota with their invasion into a stabilized precultured soil microbiota background. By comparing the growth of inoculants axenically in the same soil microcosm conditions we found that only around 1% of the potential niche available to the inoculant is free in the presence of NatCom (Fig. 3A and C). The available niche was four times greater when the inoculant was co-inoculated with NatCom than if it was introduced *after* colonization by NatCom (Fig. 3C). This was only the case for pseudomonad inoculants and not for the non-soil strain control *E. coli*, which, as expected, survived very poorly when inoculated into NatCom. Given that the starting densities of inoculant in the case of GROWING community was approximately one-tenth of the estimated Proteobacteria proportion (1×10^5^ inoculant cells g^−1^ compared to ca. 60% of 2×10^6^ cells g^−1^ soil, Fig. 1C&E) it is unlikely that these (opportunistic and fast-growing) Pseudomonas inoculants were outcompeted by faster consumption of primary growth substrates by NatCom taxa. Rather, as highlighted by random paired bead growth assays, inoculants lost productivity during growth, either by leakage of metabolites by the inoculant and their uptake by soil bacteria, by widespread secretion of growth-inhibitory substances, or both. The hypothesis of metabolite leakage and competitive loss is supported by our metatranscriptomic analysis of the specific case of inoculation of *P. veronii* and addition of toluene as a selective substrate. Our finding that, in bead confined pairs, the increased growth of soil taxa comes at the cost of decreased inoculant growth also suggests that competitive loss is a major factor in inoculant survival along with, to a lesser degree, growth inhibition.

Diversity has frequently been suggested as a key factor controlling the establishment of new species in resident microbiomes ^5,31−33^, whereas other studies have instead emphasized the importance of community productivity ^14,34^. Both factors are inherently related, given that productivity reflects and depends on community composition within the resource richness of the habitat ^35^. The more important underlying determinant of the community composition effect in this context, however, seems to be niche availability ^14−16^. Growing, habitat-adapted, taxa-diverse communities can be expected to utilize all nutrient and spatial niches, which would then limit the proliferation of incoming species (i.e., inoculants). We tested this principle directly by maintaining the same taxa-diverse NatCom in two system states: one with low (GROWING) and the other with high (STABLE) initial biomass. The GROWING NatCom expanded six times more than the STABLE community, reflective of the increased nutrient niche availability in GROWING. Three of the four tested inoculants (the three pseudomonads) indeed established better in the GROWING than the STABLE NatCom state. The effect was modest, potentially because the STABLE condition still permitted some growth of the resident community but nonetheless validates the principle. The poor proliferation in soil of *E. coli* is likely due to its general inability to exploit soil ecological niches. By providing a selective nutrient niche (i.e., toluene) within the same background we achieved two orders of magnitude higher inoculant growth, demonstrating that niche (un)availability and competition control inoculant proliferation. The concept of a unique niche for inoculant growth has been understood for infant gut succession and can be exploited by symbiotic supplements ^36,37^, whereas recent work (which employed a similar experimental system) also demonstrated how nutrient provision in the plant rhizosphere can build a specific inoculant niche ^38^. Our results show it is applicable to soil microbiota interventions.

Although niche availability explained part of the inoculant’s fates in the soil communities, we also investigated potential biological interactions, which have been considered by others as crucial for invader establishment ^5,7^. Surprisingly, we found that growth of all four inoculants was decreased in paired co-cultures with randomized soil bacteria inside micro-agarose beads (Fig. 4), whereas growth of the soil partner on average was increased. Rather than substrate competition this effect is indicative of competitive facilitation, by which the inoculants lose productivity in facilitating the growth of the partner ^39^. Thus, our findings suggest that by facilitating the growth of other soil bacteria inoculants diminish their own population expansion. We illustrated the extent of this phenomenon more clearly for the case of thriving *P. veronii* in soil exposed to toluene, which led to measurable increases of selective gene expression for aromatic compound metabolism in resident bacteria (Fig. 6). More generally, the metabolism of most bacteria results in leaking metabolites ^40^ that can become more broadly accessible to other cells in their vicinity, thereby benefiting their maintenance or growth and contributing to community diversity ^41^.

Our results demonstrated the importance of niche availability for inoculant proliferation and highlighted the consequences of facilitative metabolism on competitive outcomes. From the perspective of microbiome engineering or interventions it is important to learn the degrees of available control of a system such that intended taxonomic and/or functional changes can be achieved. This control may range from exploiting inherent and temporal available niches for growth to establishing selective (temporal) niches for one or more inoculants to thrive and exert their functionalities. Engineering soil microbiomes is particularly complicated by their inherent biotic and abiotic complexity and spatial heterogeneity, which cannot be easily tuned by process parameters like, for example, in the engineered infrastructure of a wastewater treatment plant. However, our results suggest that niche engineering is a potentially exploitable mechanism for inoculant establishment. Engineered niches need not necessarily to consist of selective carbon compounds but, potentially, could also be generated in the form of spatial niches or other limiting nutrients. *Inter alia,* our results also reflect a realistic bioremediation scenario where an inoculated bioremediation agent thrives thanks to the selective niche provided by a contaminating compound. Degradation of the contaminant can then simultaneously favor growth of other soil members, leading to the subsequent decline of the inoculant but restoration of the microbiome.

## MATERIAL AND METHODS

### Soil inoculant strains

Four strains were selected as inoculants for growth and interaction studies with soil communities: *P. veronii* 1YdBTEX2, a toluene, benzene, *m*- and *p*-xylene degrading bacterium isolated from contaminated soil ^21^; *P. putida* F1, a benzene-, ethylbenzene- and toluene-degrading bacterium from a polluted creek ^22^; *P. protegens* CHA0, a bacterium with plant-growth promoting character as a result of secondary metabolite production ^42^; and (motile) *E. coli* MG1655 (obtained from the *E. coli* Genetic stock center Yale; CGSC#8237) ^43^, as a typical non-soil dwelling bacterium. Variants of the four strains that constitutively express mCherry fluorescent protein were used. *P. veronii* 1YdBTEX2 and *P. protegens* CHA0 were tagged with mCherry (expressed under control of the P_tac_ promoter) using a mini-Tn*7* delivery system ^44^. *P. putida* F1 was tagged with the same P_tac_-mCherry cassette, cloned into and delivered by mini-Tn*5* suicide vector pBAM ^45^. *E. coli* MG1655 was tagged with the same P_tac_-mCherry cassette but on plasmid pME6012 ^46^.

### Culturing of NatCom soil microbial communities in soil microcosms

Inoculant proliferation was tested in sterile soil microcosms, with or without precolonization by resident soil microbiota. The microcosms were prepared according to the procedure described in Čaušević et al. ^25^, by complementing dried, double-sieved, and twice autoclaved silt (obtained particle size 0.5–3 mm), with soil organic solution to a final gravimetric water content of 10%. Soil organic solution was water extracted from top-soil (1–5 cm, Dorigny forest at the University of Lausanne campus). Equal volumes of soil and tap water were mixed and autoclaved for 1 h and left to cool overnight. After decanting, the liquid fraction was further centrifuged and filtered (<0.22 µm) to remove soil and plant debris. The resulting solution was autoclaved a second time to ensure complete sterility. Our soil microbiota was sourced from previously grown top soil microbial communities (NatCom) in the same type of reconstituted sterile soil material ^25^, which had been stored at room temperature (23 °C) for 1.5 years.

Soil microbiota were revived by transferring 11 g of the stored NatCom soil into 80 g sterile soil material reconstituted with 9 ml forest soil extract in a 500 ml capped Schott glass bottle (Fig. 1A, 50 replicates). Five microcosms were selected randomly for the Phase 1 analysis and the rest of the microcosm material was kept for Phase 2. The microcosms were periodically mixed on a bottle roller and incubated at room temperature (23 °C) in the dark for 28 days to allow the growth, dispersal, and colonization of the NatCom throughout the soil material. The five selected microcosms were sampled at regular intervals to assess taxa composition by amplicon sequencing and cell density by flow cytometry (see below). After 28 days, the content of all inoculated microcosms was mixed and divided into two new sets of 28 microcosms. In one set (the STABLE state) the pooled and colonized soil material (100 g) was directly transferred to new, sterilized, and empty Schott bottles without any new addition of nutrients. In the second set (the GROWING state) the colonized material (11 g) was mixed with 80 g freshly sterilized soil matrix and 9 ml forest soil extract in a new bottle. This soil-to-soil dilution allowed a new phase of active community growth.

### Introduction of inoculants in soil microcosms

For inoculation into soil microcosms all pseudomonads were grown individually from frozen glycerol stocks in Lysogeny-Broth (LB, BD Difco) supplemented with 25 µg mL^−1^ of gentamicin (30 °C) and *E. coli* was grown at 37 °C in LB with 25 µg mL^−1^ of tetracycline, to maintain the fluorescent marker. After 16 h culturing, cells were harvested by centrifugation, washed, and subsequently diluted in type 21 C minimal medium (MM; ^47^) with 0.1 mM succinate to obtain a concentration of 10^7^ cells ml^−1^. Four sets of four replicates each of STABLE or GROWING microcosms (see above) were then inoculated with either of the four strains to achieve a starting inoculant cell density of 10^5^ cells g^−1^ of soil, while one set of four remained unamended to verify sterility. Inoculants were inoculated individually (ALONE) into sterile soil microcosms (4 replicates each, same microcosm material). A final two sets of four microcosms (with or without NatCom in either the STABLE or GROWING state) were amended with *P. veronii* and toluene or with toluene alone (see below). Following inoculation the microcosms were mixed on a bottle roller and incubated at 23 °C.

### Addition of toluene to microcosms

Toluene (Fluka Analytical) was introduced to the microcosms in the gas phase via 0.5 ml pure toluene held in a heat-sealed 1-ml micropipette tip, which was placed inside a sealed 5-ml tip for additional stability, carefully placed inside the microcosms. At each mixing and sampling step, toluene tips were removed from microcosms using sterile tweezers, the level of toluene was checked and replenished to 0.5 ml, if necessary, after which the tips were replaced once the content of microcosm was mixed.

### Extraction of cells from soil microcosms

Soil community size and composition was quantified using cells washed from the soil matrix at each time point. Microcosm material (10 g) was sampled using Sterileware sampling spatulas (SP Bel-Art) and transferred to a 50 ml capped Greiner tube, after which 10 ml sterile tetrasodium-pyrophosphate decahydrate (TSP) solution (2 g l^−1^, pH 7.5, Sigma-Aldrich) was added and the mixture vortexed for 2 min at maximum speed on a Vortex-Genie 2 (Scientific Industries, Inc.). Debris was allowed to settle for 2 min and the supernatant (cell suspension) was transferred to a new tube. This suspension was then used for cell enumeration by flow cytometry or colony forming unit counting, DNA isolation, and amplicon sequencing.

### Flow cytometry cell enumeration

A portion of the cell suspension (see above) was passed through a 40 µm nylon strainer (Falcon) to remove particulate material. Two 100-μl aliquots of filtrate were then mixed with equal volumes of 4 M sodium azide solution (Sigma-Aldrich) to fix the cells. Fixed samples were kept at 4 °C until processing with flow cytometry (within 1 week). Before flow cytometry measurement, one fixed sample was stained with SYBR Green I for 15 min in the dark (Invitrogen, following manufacturer’s instructions) whereas the other remained unstained, allowing the estimation of background fluorescent particle content. Stained and non-stained suspensions (10 µl) were aspired on a CytoFLEX Flow Cytometer (Beckman Coulter) at the slow flow rate (10 µl min^−1^). Phase 1 non-inoculated microcosms were used as controls for background noise coming from soil, which was subtracted from total counts of treated microcosms. The inoculants were detected and gated based on their mCherry tag (ECD-H signal) signal in the non-SYBR Green I–stained sample series.

### Colony forming unit counting

A 100-µl aliquot of soil cell suspension was serially diluted using TSP solution and 10 µl droplets (four technical replicates) of each dilution (from 10^0^ to 10^−7^) were deposited on R2A plates (DSMZ GmbH). The plates were left to dry for 10 min and then incubated at 23 °C in the dark. Colonies were counted after 3 days of growth using a stereo microscope (Nikon SMZ25), and the corresponding community number of colony forming units (CFU) g^−1^ soil was calculated from the cell extraction procedure and its dilutions.

### Amplicon sequencing

The remaining cell suspension (9 ml) was centrifuged in a swing-out rotor (Eppendorf A-4-62 Swing Bucket Rotor) at 4000 × g for 7 min to harvest the cells. The supernatant was discarded and cell pellets were stored at –80 °C. DNA was subsequently extracted from thawed cell pellets using a DNeasy PowerSoil Pro DNA Isolation Kit (Qiagen, as per instructions by the supplier). Final yields were quantified using a Qubit dsDNA BR Assay Kit (Invitrogen), and the purified DNA solution was stored at –20 °C until library construction. Each sample (10 ng DNA input) was then used to amplify the V3–V4 variable region of the 16S rRNA gene, following the protocol by Illumina (16S Metagenomic Sequencing Library Protocol, https://support.illumina.com/documents/… documentation/chemistry_documentation/16s/16s- metagenomic-library-prep-guide-15044223b.pdf).

Samples were indexed by using the Nextera XT Index kit (v2, sets A and B, Illumina) after which the DNA was again purified, pooled, and sequenced using a MiSeq v3 paired-end protocol (Lausanne Genomic Technologies Facility). Raw reads were analyzed using the Qiime2 platform on UNIX (version 2021.8) ^48^, and amplified sequence variants (ASVs) were attributed to known taxa at 99% identity (operational taxonomic units, OTU) by comparison to the SILVA database (version 132).

### Paired inoculant-soil taxa growth assays in encapsulating agarose beads

Potential growth effects between inoculants and soil taxa were tested using random pairs of single cells encapsulated within 40–70 µm diameter polydisperse agarose beads ^24^. Inoculants were precultured as follows. *P. veronii* and *P. putida* were grown on MM agar with toluene as sole carbon source provided through the vapour phase in a desiccator, as described previously ^49^. A single colony grown after 48 h incubation at 30 °C was subsequently inoculated into 10 ml MM with 5 mM succinate as the sole carbon source and cultured for 24 h. *P. protegens* and *E. coli* colonies were cultured as described above on selective nutrient agar plates supplemented with 25 µg ml^−1^ of gentamicin or 25 µg ml^−1^ of tetracycline, respectively, and then transferred to liquid MM with 5 mM succinate. After 24 h growth, the cells were harvested from their precultures by centrifugation and resuspended in 10 ml MM. Cell suspensions were counted by flow cytometry and diluted to 2×10^7^–10^8^ cells ml^−1^ for the bead encapsulation process. Soil microorganisms were washed and purified for each encapsulation experiment from four 200 g samples of fresh soil (characteristics and location as described previously ^24^) using a similar procedure as described above for the NatComs. Purified cells were counted by flow cytometry and diluted in MM to 1×10^8^ cells ml^−1^ before encapsulation. Each of the inoculant or washed soil cell suspensions alone, or inoculant mixed in 1:1 volumetric ratio with the soil cell suspension, were then mixed with liquid low-melting agarose solution (37 °C) to produce 40–70 µm diameter agarose beads with a Poisson-average of two founder cells at start, using the procedure described previously ^24^. Per condition and type of inoculant, two batches of cell-encapsulated beads were prepared in parallel, which were pooled and then split in three replicate tubes each containing 1 ml bead solution. The encapsulation procedure produced 1.2×10^6^ beads per ml, with an estimated effective ‘bead’ volume of 10% of the total volume of the liquid phase in the incubations.

### Culture conditions for bead-encapsulated cell pairs

Three different carbon source regimes were imposed on bead-encapsulated cells: toluene, mixed carbon substrates, or soil extract. Toluene was used as an example of an inoculant-selective substrate (for *P. veronii* and *P. putida*) and was provided by partitioning from an oil phase. We diluted pure toluene 1000× in 2,2’,4,4’,6,8,8’-heptamethylnonane (Sigma Aldrich) and added 0.2 ml of this solution to each vial with 1 ml bead suspension. A further 4 ml of MM was added to the vials for the incubation. Mixed carbon substrates (Mixed-C), and soil extract were used as diverse substrates for all inoculants and soil microbes. Mixed-C solution was prepared by dissolving 16 individual compounds (Table S1) in milliQ-water (Siemens Labostar) in equimolar concentration such that the total carbon concentration of the solution reached 10 mM-C. These compounds have been used previously as soil representative substrates ^50^. In the bead incubations, the Mixed-C was diluted to 0.1 mM-C final concentration in MM (5 ml total volume per vial) to avoid excessive growth of microcolonies inside the beads, which could lead to cell escape and their proliferation outside the beads.

Soil extract for agarose beads was prepared as follows. A quantity of 100 g soil (same origin as used for the soil community cell suspension ^24^) was mixed with 200 ml 70 °C milliQ-water in a 250 ml Erlenmeyer flask and swirled on a rotatory platform for 15 min after which it was subjected to 10 min sonication in an ultrasonic bath (Telesonic AG, Switzerland). Sand particles were sedimented and the supernatant was decanted and passed through a 0.22 µm vacuum filter unit (Corning Inc.). The resulting soil extract (4 ml) was added directly to the 1 ml bead suspension in the vials. Triplicate vials per treatment and per inoculant- mixture were incubated at 25 °C with rotary shaking at 110 rpm to prevent sticking of the beads but avoid shearing damage.

### Sampling and analysis of cell growth in agarose beads

Encapsulated cell mixtures were sampled at regular time intervals (0, 6, 24, 48 and 72 h). For this, 10 µl of bead suspension was removed from the vials. Cells and microcolonies in the beads were stained with SYTO- 9 solution and imaged with epifluorescence microscopy, as described previously ^24^. Microcolony growth was quantified using a custom MATLAB (v. 2021b) image processing routine that segmented beads and microcolonies inside beads ^51^. Inoculant cell colonies were differentiated from soil cells based on having both mCherry and SYTO-9 fluorescence, whereas soil cells displayed only SYTO-9 fluorescence. Growth was calculated as the product of SYTO-9 fluorescence area and mean fluorescence intensity for each detected microcolony ^24^. Beads containing exactly one inoculant and one soil cell microcolony were selected to plot paired productivities. Productivities were compared to simulations (n = 5) of the expected paired productivity without any assumed interaction from the (subsampled) observed growth in encapsulated beads of either the inoculant or the soil cells alone. Differences among observed and expected paired productivities were evaluated from the sums across three regions as indicated in Figure 4C in an unpaired t-test.

### Metatranscriptomic analysis

To better understand the impact of adding an inoculant and/or toluene on the native soil community we took advantage of previously conducted inoculation experiments of *P. veronii* in a variety of soil types from which total RNA had been purified ^28^. These consisted of two uncontaminated soils (Clay and Silt), and one contaminated soil from a former gasification site named *Jonction*. Soils had been exposed or not to toluene and inoculated with *P. veronii*. We expected a high background of mono-aromatic compound degrading resident bacteria in *Jonction* because of its long-term contamination and, thus, more functional diversity in transcripts from aromatic-degradation pathways (Fig. S6 and S7). Soil microcosms had been sampled in an ‘early’ or a ‘late’ state (Fig. 6A) (the exact timing roughly depending on observed growth of the inoculant population)^28^. Total purified RNA from the samples was depleted for bacterial ribosomal RNAs, reverse-transcribed, indexed, and sequenced on Illumina HiSeq 2500 or NovaSeq at the Lausanne Genomic Technologies Facility following a previously described procedure ^28^.

Sequencing reads from all samples were quality controlled by BBMap (v.38.71), which removed adapters from the reads, removed reads that mapped to PhiX (a standard added to sequencing libraries) and discarded low-quality reads (trimq=14, maq=20, maxns=1, and minlength=45). Quality-controlled reads were merged using bbmerge.sh with a minimum overlap of 16 bases, resulting in merged, unmerged paired, and single reads. The reads from metatranscriptomic samples were assembled into transcripts using the SPAdes assembler ^52^ (v3.15.2) in transcriptome mode. Gene sequences were predicted using Prodigal ^53^ (v2.6.3) with the parameters -c -q -m -p meta. Gene sequences from the GenBank entry of *P. veronii* (GCA_900092355) were downloaded and clustered at 95% identity, keeping the longest sequence as representative using CD-HIT ^54^ (v4.8.1) with the parameters -c 0.95 -M 0 -G 0 -aS 0.9 -g 1 -r 1 -d 0. Gene sequences predicted from assembled transcripts were used to augment the *P. veronii* database using CD-HIT (cd-hit-est-2d -c 0.95 -M 0 -G 0 -aS 0.9 -g 1 -r 1 -d 0). Representative gene sequences were aligned against the KEGG database (release April.2022) using DIAMOND ^55^ (v2.0.15) and filtered to have a minimum query and subject coverage of 70%, requiring a bitScore of at least 50% of the maximum expected bitScore (referenced against itself).

The 145 metatranscriptome samples were then mapped to the 246,873 cluster representatives with BWA ^56^ (v0.7.17-r1188; -a), and the resulting BAM files were filtered to retain only alignments with a percentage identity of ≥95% and ≥45 bases aligned. Transcript abundance was calculated by first counting inserts from best unique alignments and then, for ambiguously mapped inserts, adding fractional counts to the respective target genes in proportion to their unique insert abundances.

### Data processing and statistical analysis

Data processing, analysis of community composition, and statistical analysis were done using GraphPad Prism (version 9.0.1) and R 4.0 (R Core Team, 2019) on RStudio (version 2022.2.3.492) using the following packages: *phyloseq* ^57^, *microbiome* ^58^, *MicrobiotaProcess*, *ggplot2* ^59^, *vegan* ^60^, *biomformat* ^61^, *tidyverse* ^62^, *reshape2* ^63^, *Biostrings* ^64^, *PMCMRplus* ^65^, *emmeans* ^66^, and *RVAideMemoire* ^67^. Chao1 values of Phase 1 samples (Day 0 to Day 23, different replicates) were compared with a Wilcoxon test. Shannon values of Phase 1 samples were compared with a Kruskal-Wallis test and by pairwise comparisons using Dunn testing with Holm’s p-value adjustment. Beta-diversity of Phase 1 community compositions (at species level) was analyzed using Unifrac distances with PCoA ordination (using *phyloseq* in R). Differences were analyzed using PERMANOVA (999 permutations) using the *adonis2* function, while data homogeneity was checked using *betadisper* function of the *vegan* package. Finally, pairwise differences between timepoints were investigated using *pairwise.perm.manova* from *RVAideMemoire* and p-values were adjusted using Holm’s method. Phase 2 Chao1 and Shannon values of GROWING and STABLE were compared using two-way repeated measures ANOVA to investigate the effect of community state and time (only significant effects are shown). For both, *post hoc* testing was done with *t* tests and p-values were adjusted with Holm’s method.

Flow cytometry data was imported using the function *fca_readfcs* ^68^ and analyzed using custom MATLAB scripts (v. 2021b). Flow cytometry counts of GROWING and STABLE NatCom were compared using two-way repeated measures ranked ANOVA. Inoculant population sizes in conditions ALONE, GROWING, or STABLE (all time points together) were compared with a Kruskal-Wallis test followed by a *post hoc* Dunn test. The same test was used to evaluate the per inoculant differences in average population size and fold-increase. Differences in percentage of inoculant survival in GROWING or STABLE conditions were tested with a one-way ANOVA followed by Tukey’s test. The effect of toluene on total community population size was examined with a Wilcoxon test (for GROWING and STABLE, separately). Changes in *P. veronii* population sizes upon introduction into toluene exposed microcosms were evaluated using Kruskal-Wallis testing with a *post hoc* Dunn test. All p-values from multiple pairwise comparisons were adjusted for multiple testing using Holm’s p-value adjustment method.

The effect of inoculant or toluene on taxa abundances was evaluated separately per replicate (randomly paired between control and treatment) and per time point using log_10_ transformed abundances. Theoretical values were inferred using a linear regression model, and taxa with abundances higher or lower than one log compared to expected value were classed as an enriched or depleted outlier, respectively. The total difference of each outlier to their expected value per condition was termed ‘total outlier distance’ and was subsequently calculated for each of the conditions (enriched and depleted taxa were quantified separately).

Microcolony productivity distributions in agarose beads were globally compared non-parametrically with the Fisher test (implemented in R 4.0) because of their non-normal nature. Productivities of paired inoculant-soil cell taxon within the same bead were evaluated by comparing to expected *null* distributions using unpaired t-tests, as described above.

The main goal of the metatranscriptomic experiments with *P. veronii* in a variety of soil types was to characterize the response of the native soil community to the addition of inoculant and/or toluene. For this, all transcripts assigned to the *P. veronii* genome were removed from the metatranscriptomic data leaving only transcripts assigned to the native soil microbial community. Length-normalized transcript abundances were then calculated by dividing the total insert counts by the length of the respective gene in kilobases. Transcript abundances per kilobase (TPK) were further converted into transcripts per kilobase million (TPM) as follows ^69^. The sum of TPK values in a sample was divided by 10^3^, and the result was used as a scaling factor for each sample. Each individual TPK value was divided by the respective scaling factor to produce the TPM values. Genes assigned to metabolic pathways associated to toluene and aromatic compound degradation were selected based on a pre-defined list of KEGG identifiers (Table S2). Representative genes for some of the highly expressed pathways were taxonomically annotated by comparing to publicly available genomes. All genes from bacterial and archaeal genomes annotated to the corresponding KEGG orthologs (K15765, K16242, K00446, K07104, K04073, K10216, K05549, K16319) in

IMG/M (Integrated microbial genomes and microbiomes: https://img.jgi.doe.gov/) were downloaded and used as a reference database to annotate all genes from the metatranscriptomics data with the same KEGG ortholog assignment. Global sequence alignment was performed with *vsearch* (v2.15) and genes were taxonomically assigned to the best hit (i.e., highest sequence identity; hits with a sequence identity below 70% were discarded). Genes assigned to ribosomal proteins were identified by a text-based query of the gene annotations. The relative abundance (proportion of TPM values) of all transcripts assigned to ribosomal proteins was used as an index of the native community growth rate. Indeed, levels of ribosomal protein transcripts have been shown previously to be well correlated with growth rate in yeast^70^, Bacteria ^71^ and Archaea ^72^ and have been proposed and used as a metric for assessing in situ growth rates from metatranscriptomic data ^73^. All statistical test results are reported in S1 Dataset.

## Data accessibility

Raw metatranscriptomic datasets of *P. veronii* inoculation into Clay, Silt, and Jonction are available from Bioproject accession number PRJNA682712, and datasets depleted from *P. veronii* reads itself can be accessed from the European Nucleotide Archive (accession numbers, ERS2210331 to ERS2210346). The raw 16S rRNA gene V3-V4 amplicon sequences for the random-paired inoculant-bead communities incubated under different substrate conditions can be accessed from the Short Read Archives under BioProject ID PRJNA661487. Finally, NatCom community profiling by 16S rRNA gene amplicon analysis is accessible through BioProject ID xxx.

## Acknowledgements

The authors thank Christoph Keel for providing us with *P. protegens* CHA0 and its tagged derivative. We are very grateful for critical reading of the manuscript by Phil Gwyther.

