## Supplementary information for "Niche Availability and Competitive Facilitation Control Proliferation of Bacterial Strains Intended for Soil Microbiome Interventions"

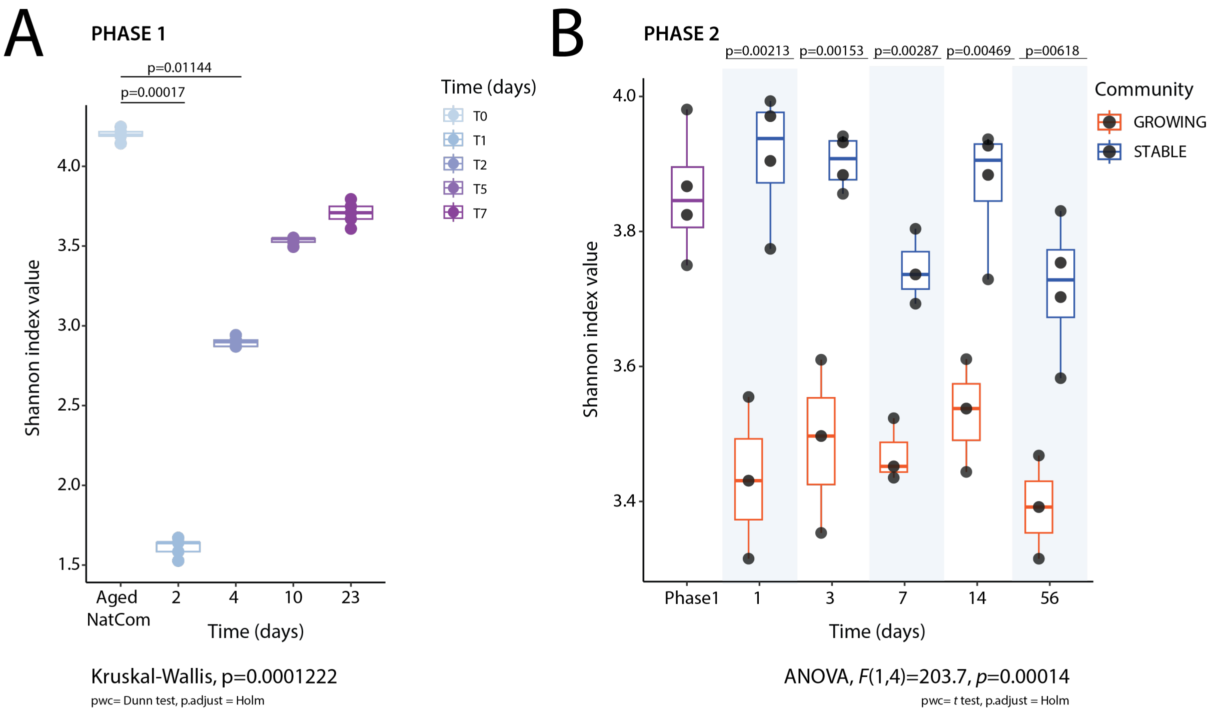


**Fig. S1. Changes in diversity of Phase 1 and Phase 2 NatComs. (A)** Shannon diversity measures calculated at amplicon sequence variant (ASV) level over time (shades from blue to magenta) for revived NatComs in Phase 1 and the aged NatCom inoculum. Dots represent individual values of five replicates. *P*-values are indicated if <0.05 after Holm's *p*-value adjustment. **(B)** Shannon diversity values for Phase 2 GROWING (orange) and STABLE (blue) NatComs over time. Phase 1 denotes the inoculum material resulting from one month of Phase 1 incubation and used as the inoculum for Phase 2 (not included in the reported statistical analysis). *P*-values above the boxplots refer to pairwise comparisons (Holm's adjustment). ANOVA refers to a repeated measures two-way ANOVA, reporting the significant effect of NatCom growth phase.

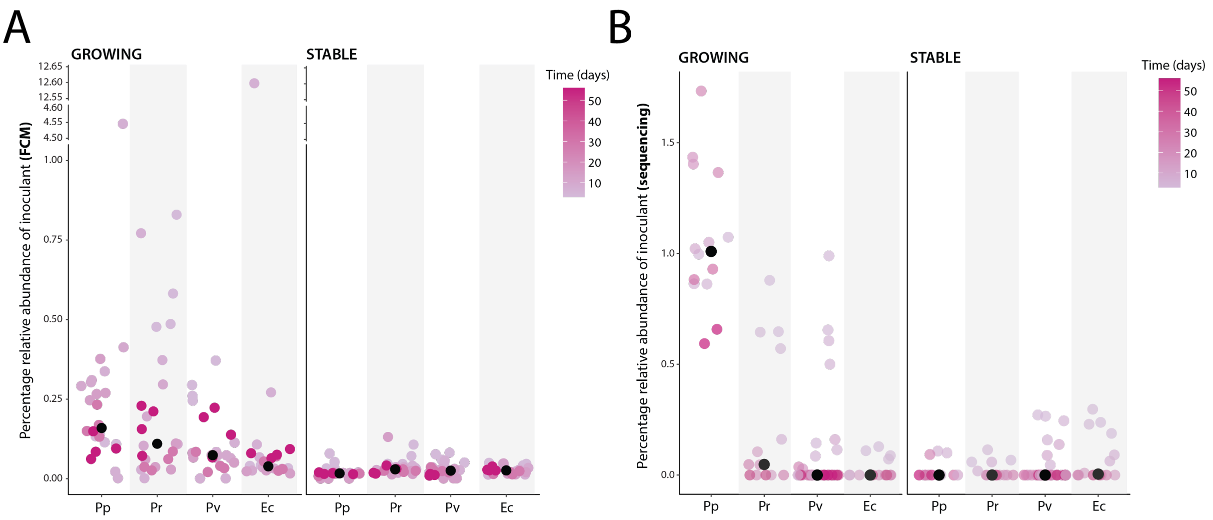


**Fig. S2. Relative abundances of inoculants in Phase 2 GROWING or STABLE NatComs. (A)** Relative abundance of each inoculant over time (time represented by the shade of magenta, according to the indicated scale), calculated by dividing the flow cytometry (FCM) counts of the inoculant (based on their mCherry fluorescent signal) by the corresponding total community counts (Syto9-stained signal) and expressed as percentage. Black dots represent the median of pooled replicates and time points per inoculant. **(B)** As in (A), with relative abundances calculated from community 16S rRNA gene amplicon composition data.

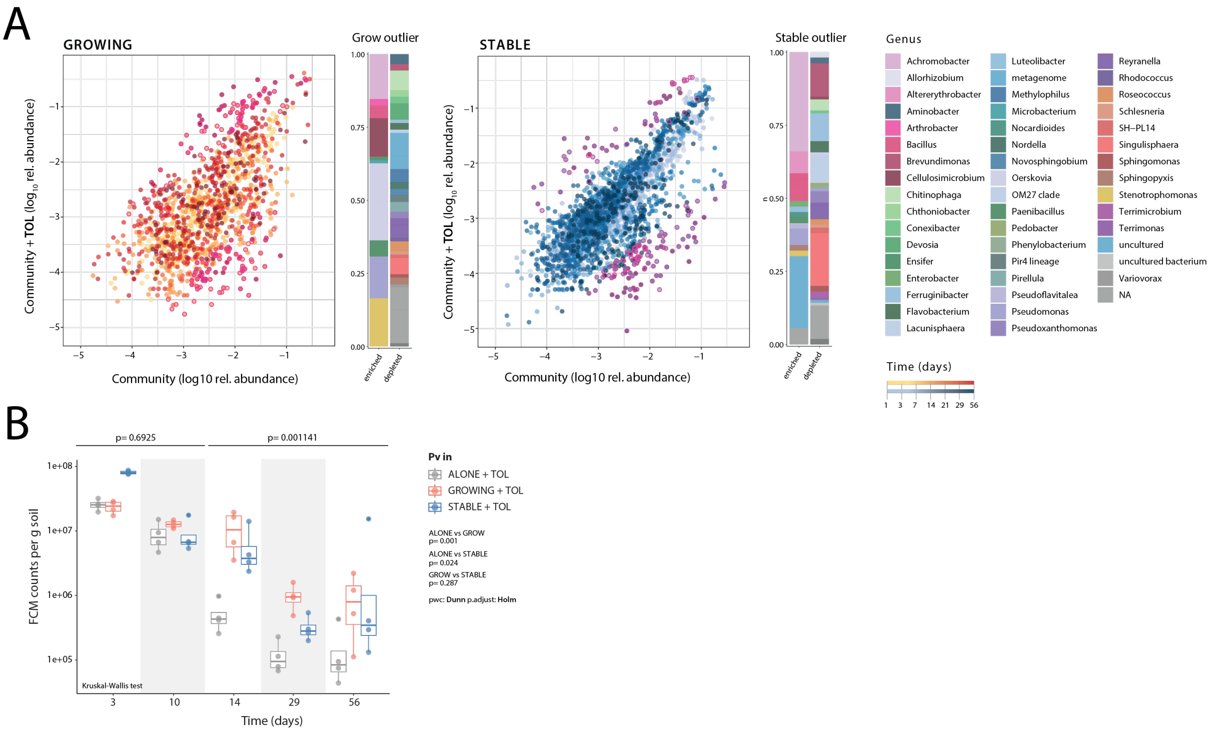


**Fig. S3. Effect of toluene addition on NatCom state and inoculant persistence. (A)** Changes of OTU abundances in GROWING and STABLE NatComs exposed or not to toluene (TOL), but without introduced *P. veronii*. Dots represent time-paired $\log_{10}$ transformed relative abundances of the same operational taxonomic units (OTU) across all biological replicates (arbitrarily paired among replicates) and treatments, as indicated. Dots with magenta circles highlight OTU abundance deviations larger than ten-fold compared to no effect (the diagonal). The composition of all enriched and depleted OTUs is shown as stacked bar subplots on the side. **(B)** *P. veronii* (Pv) population sizes (FCM counts of mCherry fluorescence) in microcosms exposed to toluene either ALONE or in presence of GROWING or STABLE resident community. *P*-values above plots correspond to Kruskal-Wallis comparisons, grouped per 'early' (Day 3 and 10) and 'late' time points (Day 14, 29, and 56). *P*-values on the side correspond to Dunn pairwise comparisons for the late phase sizes of the *P. veronii* populations.

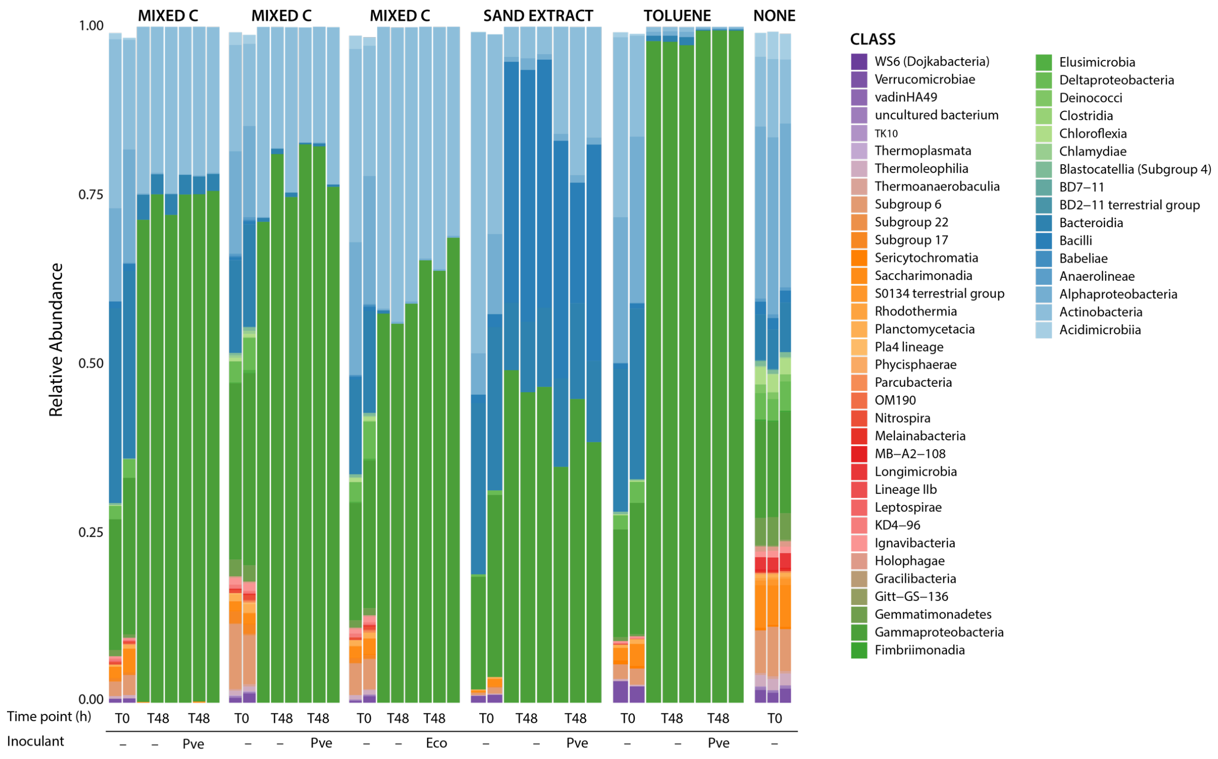


**Fig. S4. Taxa diversity in encapsulated soil microbiota-inoculant mixtures.** Stacked bars of class-level relative abundances of attributed OTUs from 16S rRNA gene amplicon sequences for washed soil microbiota on six independent occasions before encapsulation (T0, – inoculant; in duplicate) or after encapsulation and 48 h incubation in absence (T48, –) or presence of inoculant (T48, Pve = *P. veronii*; Eco = *E. coli*). Medium conditions are indicated at the top of the panel (mixed C = mixture of 16 carbon sources, sand extract = washed autoclaved organic carbon from soil, toluene = addition of toluene dissolved in heptamethylnonane, and none = medium only without carbon source).

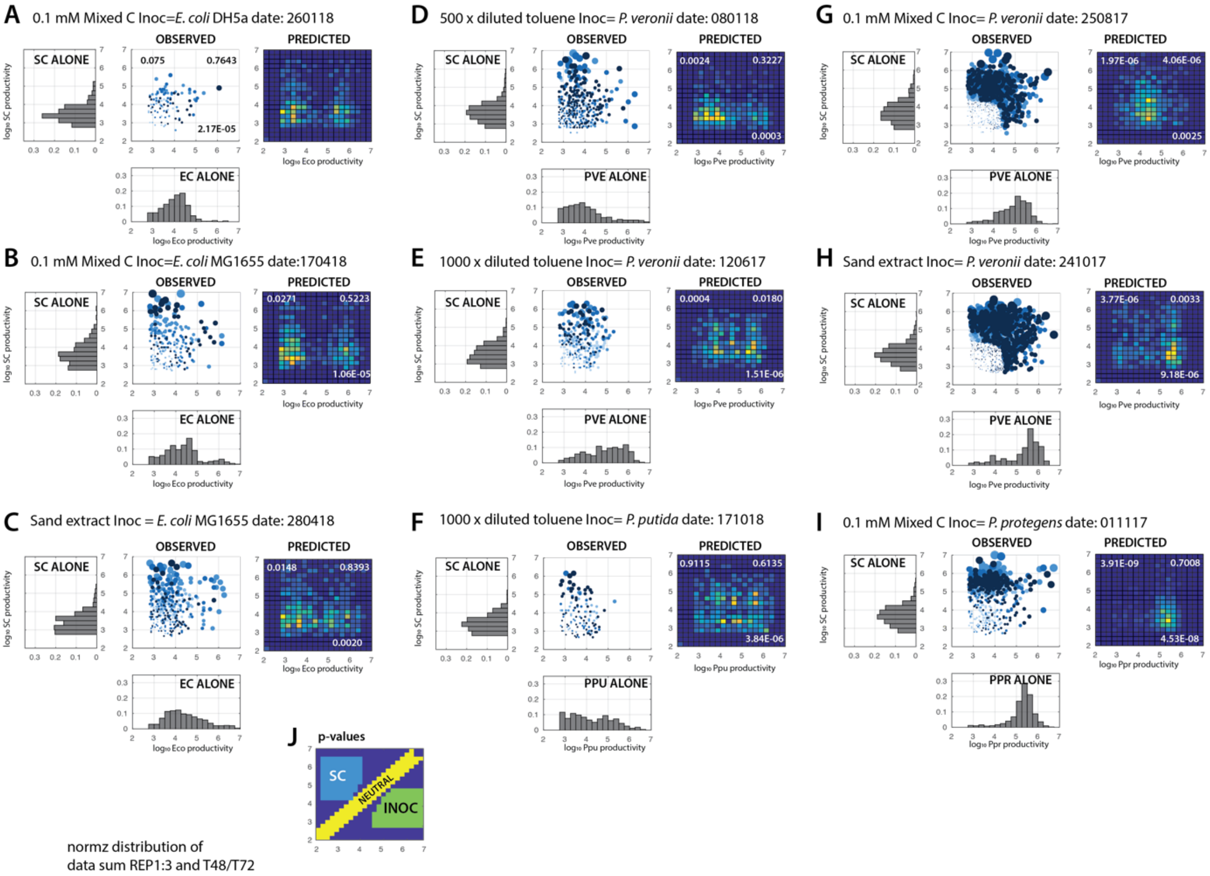


**Fig. S5. Expected versus observed paired productivities of inoculants with random resident soil cells. (A) – (I)** Paired productivity plot of beads with only a single inoculant and a single soil cell microcolony (OBSERVED) versus beads with inoculant or soil cells alone (summed from time points 24, 48, and 72 h). Circles are proportional to the sum of the measured microcolony sizes (light to dark blue represent time points 0, 6, 24, 48, and 72 h). The heatmap (PREDICTED) shows the expected paired bead summed productivities from the individual measured microcolony sizes (i.e., inoculant and SC ALONE) for the same number of beads as analyzed by microscopy, with yellow colors corresponding to high incidence and dark blue to low. *P*-values correspond to the two-tailed t-test comparison of the variation of the total measured paired productivities inside the three regions ($n$ = 3; 24, 48, and 72 h) to that in the simulations ($n$ = 5), as indicated in panel J. The upper left region shows higher SC productivity than expected, the lower right regionshows lower inoculant productivity than expected, and the diagonal shows thesame productivity for both microcolonies in a pair (EC =*E. coli*, PVE =*P. veronii*, PPU = *P. putida*, and PPR = *P. protegens*).

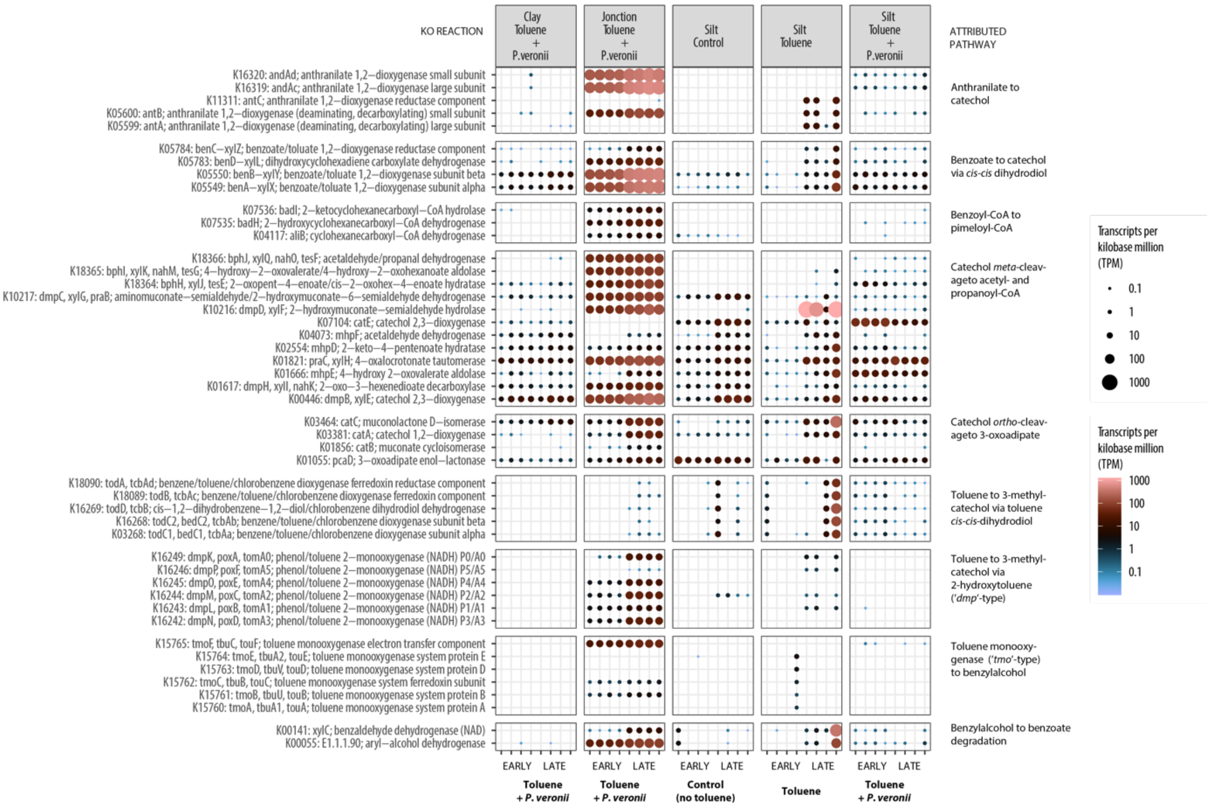


**Fig. S6. Enrichment of toluene degradation pathways in soil microbiota in presence or absence of added *P. veronii*.** Plots show assigned transcript abundances (in TPM (transcripts per kilobase million) as circle sizes and colors according to scales on the right) in the various sample incubations for KEGG orthology reactions (as specified on the left of the panels and grouped per pathway on the right). TPM values of transcripts annotated to the aromatic compound metabolic steps on the right for the different soils, conditions, and timepoint (early or late, see panel A of Fig. 6 in the main text). Data points show individual values from quadruplicate experiments.

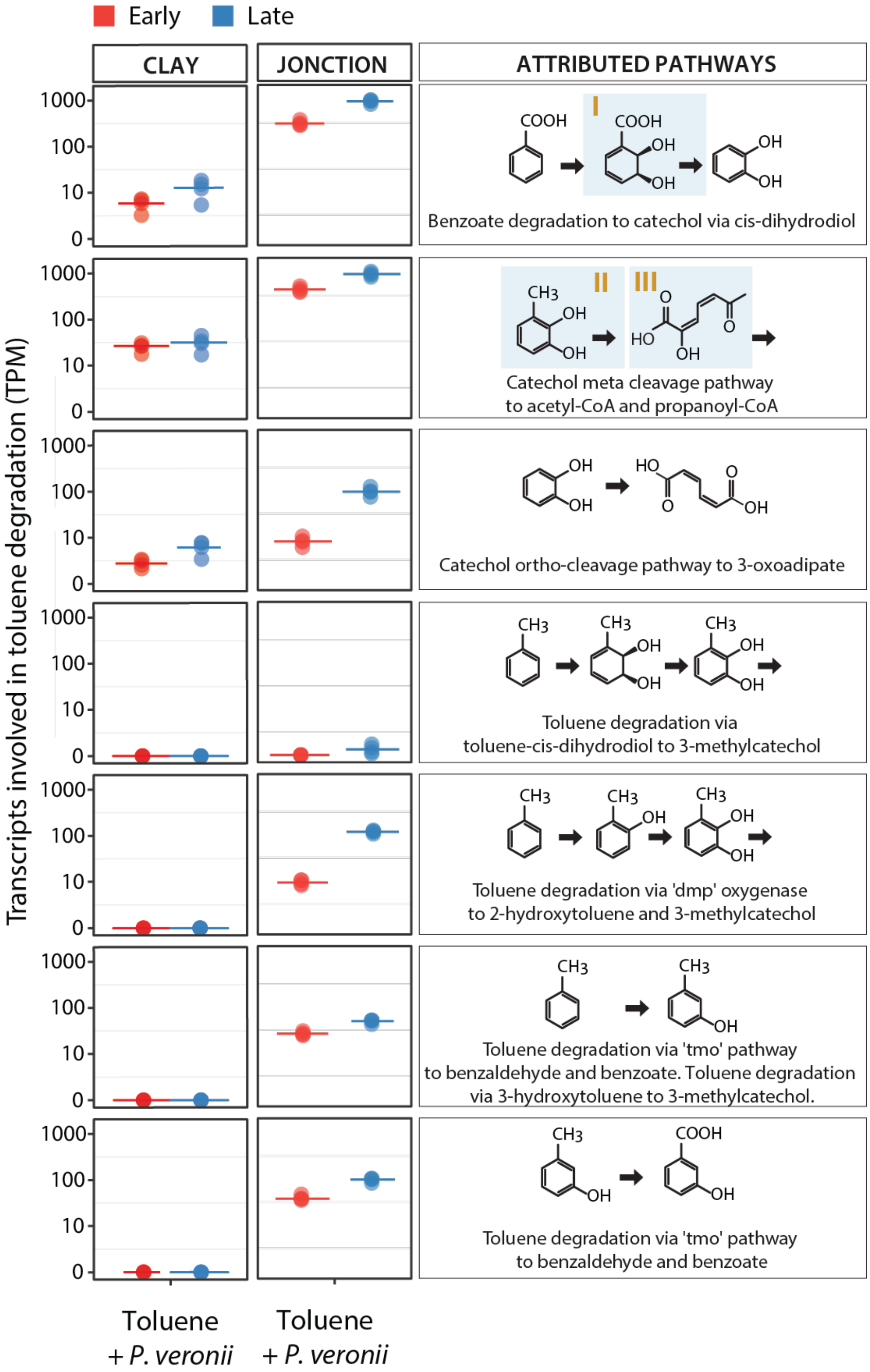


**Fig. S7. Exploitation of toluene degradation products from *P. veronii* by microbiota in Clay and Jonction**. TPM values of transcripts annotated to the aromatic compound metabolic steps on the right for Clay and Jonction soils inoculated with *P. veronii* and tested at two timepoints (early or late, Fig. 6A). Data points show individual values from quadruplicate experiments, presented as dots, and a line indicating their median value.

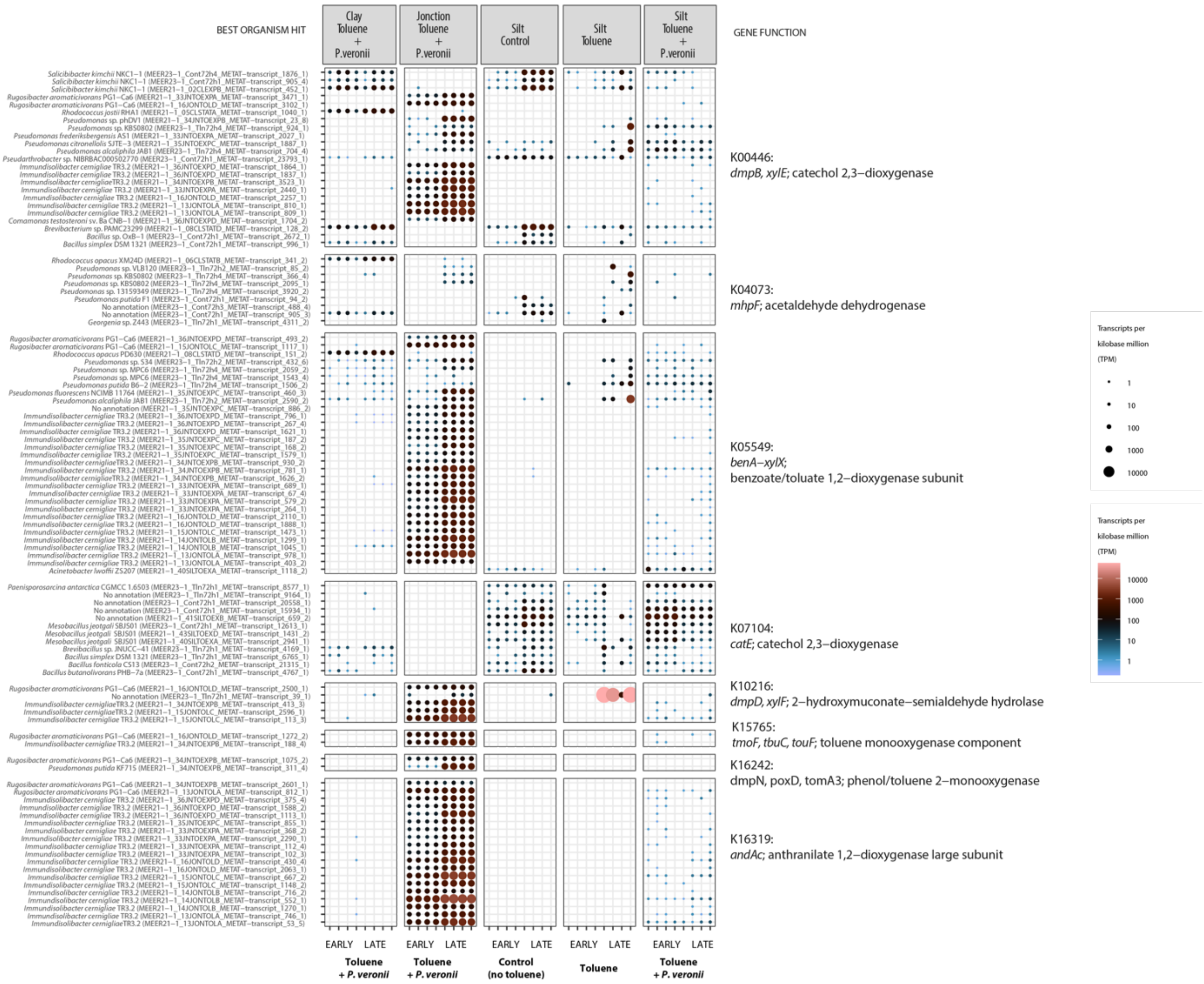


**Fig. S8. Taxa attribution of enriched toluene degradation pathways among soil microbiota in the presence or absence of added *P. veronii*.** Plots show assigned transcript abundances (in TPM as circle sizes and colors according to scales on the right) for specific KEGG orthology reactions and representative genes (as specified on the right of the panels) attributed to the best-hit taxa score (on the left). TPM values of transcripts for the different soils, conditions, and timepoints (early or late, see panel A of Fig. 6 in the main text). Data points show individual values from quadruplicate experiments.

**Table S1: Composition of 16 carbon source minimal medium for soil bacteria growth**.

| | | Mw | No C-atoms | Mass to weigh (mg per L) |
|---|---|---|---|---|
| 1 | L-Arginine | 174.2 | 6 | 13.10 |
| 2 | D-Xylose | 150.1 | 6 | 11.29 |
| 3 | L-Aspartic acid potassium salt | 209.3 | 4 | 15.74 |
| 4 | 4-Hydroxybenzoic acid | 144.0 | 7 | 10.83 |
| 5 | L-Serine | 105.1 | 3 | 7.90 |
| 6 | beta-Hydroxy Butyric Acid | 104.1 | 4 | 7.83 |
| 7 | D-Cellobiose | 342.3 | 12 | 25.74 |
| 8 | alpha-D-Lactose | 360.3 | 12 | 27.09 |
| 9 | Putrescine | 88.15 | 4 | 6.63 |
| 10 | Itaconic acid | 130.1 | 5 | 9.78 |
| 11 | Alpha-D-glucose-1-phosphate | 304.1 | 6 | 22.86 |
| 12 | N-acetyl-D-glucosamine | 221.2 | 8 | 16.63 |
| 13 | D-Mannitol | 182.2 | 6 | 13.70 |
| 14 | Meso-erythitol | 112.2 | 4 | 8.44 |
| 15 | Galacturonic acid | 194.1 | 6 | 14.59 |
| 16 | Tween 20 | 604.8 | 40 | 45.47 |
| | Total | | 133 C | |

Final concentration for C = 0.1 mM
Equivalent for 10 mM C on proportion of C-atoms in the compound (e.g., 6/133*10=0.451)
Conversion factor, e.g., 0.451/6=0.07519 mM of the compound to be added
Mass to weigh: conversion factor x Mw, e.g., 0.07519 * 174.2 = 13.10 mg per L

**Table S2. KEGG orthologs used as metabolic markers for toluene and aromatic compound degradation.** List of KEGG orthologs (KO terms) manually extracted and curated to characterize different known pathways for toluene and aromatic compound degradation. Where possible, each KO term has been assigned to a single pathway; otherwise a KO term might be indicative of more than one pathway.

| KO | Attributed process | Attributed pathway |
|---|---|---|
| **K07540** | Toluene degradation | Toluene degradation via benzylsuccinate to benzoyl-CoA: (anaerobic) |
| **K07543** | Toluene degradation | Toluene degradation via benzylsuccinate to benzoyl-CoA: (anaerobic) |
| **K07544** | Toluene degradation | Toluene degradation via benzylsuccinate to benzoyl-CoA: (anaerobic) |
| **K07545** | Toluene degradation | Toluene degradation via benzylsuccinate to benzoyl-CoA: (anaerobic) |
| **K07547** | Toluene degradation | Toluene degradation via benzylsuccinate to benzoyl-CoA: (anaerobic) |
| **K07548** | Toluene degradation | Toluene degradation via benzylsuccinate to benzoyl-CoA: (anaerobic) |
| **K05749** | Toluene degradation | Toluene degradation via benzylsuccinate to benzoyl-CoA: (anaerobic) |
| **K07550** | Toluene degradation | Toluene degradation via benzylsuccinate to benzoyl-CoA: (anaerobic) |
| **K15760** | Toluene degradation | Toluene degradation via 'tmo' pathway to benzaldehyde and benzoate / Toluene degradation via 3-hydroxytoluene to 3-methylcatechol |
| **K15761** | Toluene degradation | Toluene degradation via 'tmo' pathway to benzaldehyde and benzoate / Toluene degradation via 3-hydroxytoluene to 3-methylcatechol |
| **K15762** | Toluene degradation | Toluene degradation via 'tmo' pathway to benzaldehyde and benzoate / Toluene degradation via 3-hydroxytoluene to 3-methylcatechol |
| **K15763** | Toluene degradation | Toluene degradation via 'tmo' pathway to benzaldehyde and benzoate / Toluene degradation via 3-hydroxytoluene to 3-methylcatechol |
| **K15764** | Toluene degradation | Toluene degradation via 'tmo' pathway to benzaldehyde and benzoate / Toluene degradation via 3-hydroxytoluene to 3-methylcatechol |
| **K15765** | Toluene degradation | Toluene degradation via 'tmo' pathway to benzaldehyde and benzoate / Toluene degradation via 3-hydroxytoluene to 3-methylcatechol |
| **K00055** | Toluene degradation | Toluene degradation via 'tmo' pathway to benzaldehyde and benzoate |
| **K00141** | Toluene degradation | Toluene degradation via 'tmo' pathway to benzaldehyde and benzoate |
| **K03380** | Toluene degradation | Toluene degradation via 3-hydroxytoluene to 3-methylcatechol |
| **K03268** | Toluene degradation | Toluene degradation via toluene-cis-dihydrodiol to 3-methylcatechol |
| **K16268** | Toluene degradation | Toluene degradation via toluene-cis-dihydrodiol to 3-methylcatechol |
| **K18089** | Toluene degradation | Toluene degradation via toluene-cis-dihydrodiol to 3-methylcatechol |
| **K18090** | Toluene degradation | Toluene degradation via toluene-cis-dihydrodiol to 3-methylcatechol |
| **K16269** | Toluene degradation | Toluene degradation via toluene-cis-dihydrodiol to 3-methylcatechol |
| **K16249** | Toluene degradation | Toluene degradation via 'dmp' oxygenase to 2-hydroxytoluene and 3-methylcatechol |
| **K16243** | Toluene degradation | Toluene degradation via 'dmp' oxygenase to 2-hydroxytoluene and 3-methylcatechol |
| **K16244** | Toluene degradation | Toluene degradation via 'dmp' oxygenase to 2-hydroxytoluene and 3-methylcatechol |
| **K16242** | Toluene degradation | Toluene degradation via 'dmp' oxygenase to 2-hydroxytoluene and 3-methylcatechol |
| **K16245** | Toluene degradation | Toluene degradation via 'dmp' oxygenase to 2-hydroxytoluene and 3-methylcatechol |

| | | |
|---|---|---|
| **K16246** | Toluene degradation | Toluene degradation via 'dmp' oxygenase to 2-hydroxytoluene and 3-methylcatechol |
| **K05549** | Benzoate degradation | Benzoate degradation to catechol via cis-dihydrodiol |
| **K05550** | Benzoate degradation | Benzoate degradation to catechol via cis-dihydrodiol |
| **K05784** | Benzoate degradation | Benzoate degradation to catechol via cis-dihydrodiol |
| **K05783** | Benzoate degradation | Benzoate degradation to catechol via cis-dihydrodiol |
| **K03381** | Catechol cleavage | Catechol ortho cleavage pathway to 3-oxoadipate |
| **K01856** | Catechol cleavage | Catechol ortho cleavage pathway to 3-oxoadipate |
| **K03464** | Catechol cleavage | Catechol ortho cleavage pathway to 3-oxoadipate |
| **K01055** | Catechol cleavage | Catechol ortho cleavage pathway to 3-oxoadipate |
| **K14727** | Catechol cleavage | Catechol ortho cleavage pathway to 3-oxoadipate |
| **K00446** | Catechol cleavage | Catechol meta cleavage pathway to acetyl-CoA and propanoyl-CoA |
| **K07104** | Catechol cleavage | Catechol meta cleavage pathway to acetyl-CoA and propanoyl-CoA |
| **K10217** | Catechol cleavage | Catechol meta cleavage pathway to acetyl-CoA and propanoyl-CoA |
| **K01821** | Catechol cleavage | Catechol meta cleavage pathway to acetyl-CoA and propanoyl-CoA |
| **K01617** | Catechol cleavage | Catechol meta cleavage pathway to acetyl-CoA and propanoyl-CoA |
| **K10216** | Catechol cleavage | Catechol meta cleavage pathway to acetyl-CoA and propanoyl-CoA |
| **K18364** | Catechol cleavage | Catechol meta cleavage pathway to acetyl-CoA and propanoyl-CoA |
| **K02554** | Catechol cleavage | Catechol meta cleavage pathway to acetyl-CoA and propanoyl-CoA |
| **K18365** | Catechol cleavage | Catechol meta cleavage pathway to acetyl-CoA and propanoyl-CoA |
| **K01666** | Catechol cleavage | Catechol meta cleavage pathway to acetyl-CoA and propanoyl-CoA |
| **K18366** | Catechol cleavage | Catechol meta cleavage pathway to acetyl-CoA and propanoyl-CoA |
| **K04073** | Catechol cleavage | Catechol meta cleavage pathway to acetyl-CoA and propanoyl-CoA |
| **K04112** | Benzoyl-CoA degradation | Benzoyl-CoA degradation, benzoyl-CoA => 3-hydroxypimeloyl-CoA |
| **K04113** | Benzoyl-CoA degradation | Benzoyl-CoA degradation, benzoyl-CoA => 3-hydroxypimeloyl-CoA |
| **K04114** | Benzoyl-CoA degradation | Benzoyl-CoA degradation, benzoyl-CoA => 3-hydroxypimeloyl-CoA |
| **K04115** | Benzoyl-CoA degradation | Benzoyl-CoA degradation, benzoyl-CoA => 3-hydroxypimeloyl-CoA |
| **K19515** | Benzoyl-CoA degradation | Benzoyl-CoA degradation, benzoyl-CoA => 3-hydroxypimeloyl-CoA |
| **K19516** | Benzoyl-CoA degradation | Benzoyl-CoA degradation, benzoyl-CoA => 3-hydroxypimeloyl-CoA |
| **K07537** | Benzoyl-CoA degradation | Benzoyl-CoA degradation, benzoyl-CoA => 3-hydroxypimeloyl-CoA |
| **K07538** | Benzoyl-CoA degradation | Benzoyl-CoA degradation, benzoyl-CoA => 3-hydroxypimeloyl-CoA |
| **K07539** | Benzoyl-CoA degradation | Benzoyl-CoA degradation, benzoyl-CoA => 3-hydroxypimeloyl-CoA |
| **K04116** | Benzoyl-CoA degradation | Benzoate degradation, cyclohexanecarboxylic acid =>pimeloyl-CoA |
| **K04117** | Benzoyl-CoA degradation | Benzoate degradation, cyclohexanecarboxylic acid =>pimeloyl-CoA |
| **K07534** | Benzoyl-CoA degradation | Benzoate degradation, cyclohexanecarboxylic acid =>pimeloyl-CoA |
| **K07535** | Benzoyl-CoA degradation | Benzoate degradation, cyclohexanecarboxylic acid =>pimeloyl-CoA |
| **K07536** | Benzoyl-CoA degradation | Benzoate degradation, cyclohexanecarboxylic acid =>pimeloyl-CoA |
| **K05599** | Anthranilate degradation | Anthranilate degradation to catechol |
| **K05600** | Anthranilate degradation | Anthranilate degradation to catechol |
| **K11311** | Anthranilate degradation | Anthranilate degradation to catechol |
| **K16319** | Anthranilate degradation | Anthranilate degradation to catechol |
| **K16320** | Anthranilate degradation | Anthranilate degradation to catechol |
| **K18248** | Anthranilate degradation | Anthranilate degradation to catechol |
| **K18249** | Anthranilate degradation | Anthranilate degradation to catechol |
